# Charting a narrow course for direct electron uptake-facilitated electromicrobial production

**DOI:** 10.1101/2022.05.28.493842

**Authors:** Anthony J. Abel, Jeremy D. Adams, Jacob M. Hilzinger, Adam P. Arkin

**Affiliations:** Department of Chemical and Biomolecular Engineering, University of California, Berkeley, CA 94720, USA; Department of Bioengineering, University of California, Berkeley, CA 94720, USA; Environmental Genomics and Systems Biology Division, Lawrence Berkeley National Laboratory, 1 Cyclotron Road, Berkeley, CA 94720, USA

**Author notes:** Correspondence should be addressed to A.P.A.

## Abstract

Electromicrobial production (EMP) processes based on CO_2_-fixing microbes that directly accept electrons from a cathode have received significant attention in the past decade. However, fundamental questions about the performance limits and viability of this strategy remain unanswered. Here, we sought to determine what would be necessary for such a system to compete with alternative sustainable production technologies based on H_2_-mediated EMP and traditional bioprocessing with crop feedstocks. Using global warming potential as the metric for comparison, we show that each EMP process can outperform sugarcane-based sucrose production. Following a stoichiometric and energetic analysis, direct electron uptake-based EMP would need to achieve a current density >48 mA/cm^2^ to reach parity with the H_2_- mediated system. Because this is currently only practical with a gas diffusion electrode (GDE) architecture, we developed a physical model of the proposed bio-GDE and used it to determine the conditions that a microbial catalyst would experience in a reactor. Our analysis demonstrates that unavoidable inefficiencies in the reactor (e.g., kinetic overpotentials and Ohmic losses) require additional energy input, increasing the breakeven current density to ∼91 mA/cm^2^. At this current density, the microbial catalyst would need to withstand a pH >10.4 and a total salinity >18.8%. Because currently-known electroautotrophs are not adapted to such extreme conditions, we discuss potential improvements to reactor design that may alleviate these challenges, and consider the implications these results have on the engineerability and feasibility of direct electron uptake-based EMP.

## Introduction

Electromicrobial production (EMP) has received significant attention as a sustainable alternative to fossil fuel-based commodity chemical production.^1–5^ In this strategy, electricity or electrochemically-derived mediator molecules act as the primary energy source for microbial transformation of CO_2_ into value-added products. EMP processes encompass a wide variety of systems based on both mediated and direct transfer of reducing power to microorganisms. In mediated systems, an electrochemically-reduced molecule, such as hydrogen (H_2_), carbon monoxide (CO), or formate (HCOO^-^) is fed to microbes as an energy source driving microbial growth and product formation.^6–9^ In direct systems, electrons are passed directly into cellular energy pools (*e.g*., the quinone pool or NADH pool) by electron conduit proteins that traverse the cell wall and/or outer membrane and make direct electrical contact with the cathode.^10–12^ This latter strategy, direct electron uptake-based EMP (dEMP), has generated significant excitement because it avoids relying on the transfer of sparingly soluble gases (H_2_, CO) into liquid phases for microbial growth (as in some mediated systems), it may obviate expensive and rare catalyst materials that are typically used in electrochemistry, and it can reduce the overall balance-of-systems intensity by integrating multiple functionalities into the same system.

To realize this promise, dEMP systems must meet several design criteria simultaneously. First, microbes must be robustly attached to a conductive cathode material for transfer of electrons into protein electron conduits with high Faradaic and energetic efficiency. Second, the amount of biomass must be sufficient to achieve a high current density,^13^ but this thick “biofilm” must minimize electronic resistivity and enable the rapid transport of key substrates like CO_2_ and any necessary nutrients.^14,15^ Third, the microorganism must efficiently use the supplied energy to generate a product of interest from CO_2_ with minimal diversion to cellular maintenance, biomass, or byproducts.^15^ Finally, the microorganism must thrive in the electrochemical environment, which is likely to be highly alkaline and saline at high current densities.^15,16^

Perhaps due to the challenges of these constraints, progress towards practical dEMP systems has remained limited. The original systems relied on acetogens^17^ and methanogens^18^ to fix CO_2_, although it was later determined that much, if not all, of the reducing power supplied to these organisms was mediated by electrochemical H_2_ generation.^19,20^ Recently, direct electron transfer mechanisms in some methanogens have been confirmed,^21^ but a crucial feature of acetogens and methanogens is that a large fraction of the fixed carbon is diverted to acetate or methane as a consequence of their metabolism.^22,23^ Hence, even if genetic tools are rapidly developed for these organisms, high selectivity towards an arbitrary product in a single reactor is unlikely.

Product spectrum and selectivity problems may be addressed by recent discoveries of a significant number of alternative electroautotrophs.^14,24–26^ Perhaps most notably, Bose and coworkers have engineered *Rhodopseudomonas palustris* to produce the bioplastic *poly*-hydroxybutyrate^27^ and the biofuel *n*-butanol^28^ during cathode-associated growth. In addition, we have recently developed a phylogenetic-driven discovery pipeline for putative electroautotrophs, which identified >70 potential electroautotrophic organisms.^14^

Proteins involved in extracellular electron transfer are also increasingly well-characterized,^10,29^ suggesting that strategies to engineer electroautotrophy in suitable microbial chassis may be possible in the near future. However, known and characterized electron conduits are not exhaustive of the myriad ecological mechanisms for electron transfer.^10,11,29^ For example, the most well-studied electron conduits (*e.g*., the Mtr proteins in *Shewanella* spp.) interface with the quinone pool in the inner membrane.^30–33^ Electron uptake based on this mechanism cannot support acetogenesis-, methanogenesis-, and sulfate reduction-based metabolisms because these terminal electron acceptors are thermodynamically uphill of the quinone pool. While acetogens probably rely on H_2_-mediated electron uptake,^20^ this suggests that methanogens and sulfate reducing microbes use some strategy for direct electron uptake that interfaces directly with the NAD^+^/NADH pool, a hypothesis that is supported by the redox potential of electron uptake in these organisms,^21,34^ although the specific mechanisms remain unclear. Hence, this conduit would need to be functionally and genetically characterized prior to its reconstitution in a suitable host organism for dEMP applications.

Recently, uncertainty has grown about the promise of dEMP.^16,35^ Mediated systems continue to make gains in scale-up,^36–38^ product spectrum diversity,^39–41^ and process modelling;^5,9^ they rely on well-characterized metabolic pathways (*i.e*., those of Knallgas, formatotrophic, or aceto-/methanogenic microbes); and, in the case of H_2_- and CO-mediated systems, they avoid challenges associated with incompatible electrolyte/medium requirements by transferring reducing power through the gas phase.^42^ Moreover, questions about the pH and salt tolerance, in addition to the productive capacity of cathodic biofilms, have also raised serious questions about the viability of dEMP.^13,16,35^

Here, we take a systematic approach to determine the feasibility of dEMP processes. We use a life cycle assessment (LCA) framework to quantify the carbon footprint (global warming potential) of this process in comparison to a well-established alternative EMP strategy, H_2_-mediated production, and a legacy process, sugarcane growth. We follow the LCA standards set by ISO 14040^43^ and 14044^44^ to quantify greenhouse gasses (in kg CO_2_-equivalents) emitted during the production of all materials and the generation of all energy necessary for a given process. We use sucrose as an example product in our analysis for three reasons. First, sucrose has a well-defined and portable secretion mechanism,^39,45^ which is necessary for direct electron uptake-based systems since microbes are immobilized on the cathode material. Second, sucrose is a useful substrate for downstream production of a wide variety of biomolecules via workhorse heterotrophic microbes such as *E. coli* and *S. cerevisiae*. Third, sucrose production from traditional crops (*e.g*., sugarcane) has well-defined life cycle impacts,^46^ which provides a solid benchmark for comparison with the EMP processes modeled here.

We began by calculating the metabolic efficiency of carbon fixation to sucrose via direct electron uptake relying on several possible physiological mechanisms for electron uptake, microbial respiration, and carbon fixation (Fig. 1a). Next, we developed a process model for the EMP scheme that, together with our energy demand calculations, enabled us to calculate the global warming potential (GWP) associated with sucrose production for the direct electron uptake-based system (Fig. 1b). Finally, we developed a multiphysics model of a gas diffusion bioelectrode (bioGDE) architecture (Fig. 1c) that could overcome the mass transfer limits of standard dEMP reactor designs and allowed us to calculate both the inefficiencies inherent to the direct electron uptake system as well as the relevant operating conditions (pH and salinity) that microbes would experience in such a reactor.

**Figure 1.**
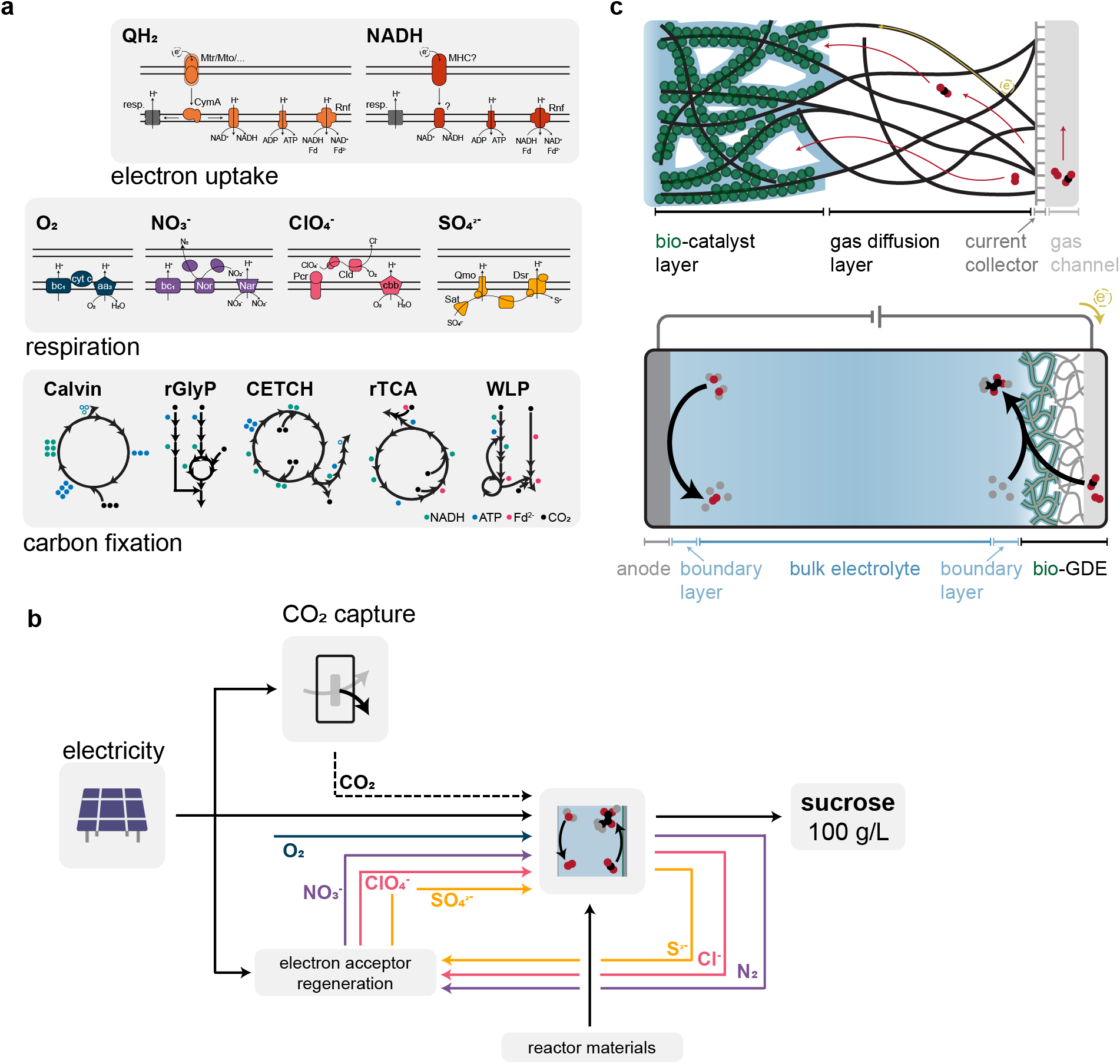
Overview of direct electron uptake-based metabolism and processes. (**a**) The three metabolic modules (electron uptake, respiration, and carbon fixation) necessary to support electroautotrophic growth and biochemical production. NADH, ATP, and ferredoxin (Fd^2-^) demands for each carbon fixation pathway are shown for pyruvate as the end product. (**b**) Schematic of the modeled electromicrobial production process. (**c**) Diagram of the modeled gas diffusion bioelectrode (bio-GDE) and the CO_2_-electrolyzer using a bio-GDE to fix carbon. Distances in (**c**) are not to scale.

Our results demonstrate that, at the current density the dEMP reactor would need to operate at to break even with an H_2_ mediated system (∼91 mA/cm^2^), the microbial catalyst would experience an average pH of ∼10.45 and an average salinity of ∼18.8%. Although some microbes exist that can tolerate such extreme conditions, enabling electroautotrophy would require complex metabolic rewiring in organisms with limited genetic tractability, and engineering extreme pH and salinity tolerance into currently-known electroactive microbes may also prove challenging. We therefore evaluate reactor improvements that could alleviate pH and salinity extremes experienced by the microbe. In contrast to the substantial hurdles facing dEMP systems, H_2_-mediated sucrose production has been demonstrated already (albeit at a modest carbon efficiency),^39^ and strategies to enhance the product yield can rely on insight gained from decades of equivalent study in cyanobacteria.^45^ Hence, our comprehensive analysis charts a narrow course for the viability of dEMP systems.

## Results and Discussion

### Physiology, stoichiometry, and energy efficiency

We first sought to determine the metabolic efficiency of direct electron uptake-facilitated carbon fixation to sucrose. We employed a stoichiometric and energetic analysis based on the three metabolic modules necessary for electroautotrophy (Fig. 1a) as described in detail in the Supplementary Information. Briefly, we determined the ATP, NAD(P)H, and reduced ferredoxin (Fd^2-^) demand for sucrose production for each of five natural or synthetic carbon fixation pathways (CFPs): the Calvin cycle,^47^ the reductive glycine pathway (rGlyP),^48,49^ the crotonyl-CoA/ethylmalonyl-CoA/hydroxybutyryl-CoA (CETCH) cycle,^50^ the reductive tricarboxylic acid (rTCA) cycle,^51^ and the Wood-Ljungdahl pathway (WLP).^52^ Next, we calculated the stoichiometry of ATP, NAD(P)H, and Fd^2-^ regeneration via respiration using oxygen (O_2_), nitrate (NO_3_^-^), perchlorate (ClO_4_^-^), and sulfate (SO_4_^2-^) as terminal electron acceptors. We used QH_2_ and NADH as the original source of electrons for these calculations based on two possible electron transfer mechanisms: electron deposition into the quinone pool, and electron deposition into the NADH pool. Finally, we determined the electron demand (mol e^-^ per mol sucrose) required to support energy carrier regeneration. The results of this analysis are compiled in Table 1.

**Table 1.** Electron demand, energetic efficiency, and energy demand for sucrose production.

| Terminal electron acceptor | Electron sink | Carbon fixation pathway | Electron demand ($\text{mol e}^-/\text{mol}$)* | Energy efficiency (%) | Energy demand ($\text{kWh/kg}$)* |
| --- | --- | --- | --- | --- | --- |
| $\text{O}_2$ | $\text{QH}_2$ | Calvin | 121 | 55 | 8.70 |
|  |  | rGlyP | 101 | 66 | 7.26 |
|  |  | CETCH† | 120 | 55 | 8.60 |
|  |  | rTCA‡ | -- | -- | -- |
|  |  | WLP‡ | -- | -- | -- |
|  | NADH | Calvin | 75.3 | 69 | 6.89 |
|  |  | rGlyP | 62.0 | 84 | 5.67 |
|  |  | CETCH<sup>†</sup> | 72.7 | 72 | 6.65 |
|  |  | rTCA<sup>‡</sup> | -- | -- | -- |
|  |  | WLP<sup>‡</sup> | -- | -- | -- |
| NO<sub>3</sub><sup>-</sup> | QH<sub>2</sub> | Calvin | 322 | 21 | 23.1 |
|  |  | rGlyP | 247 | 27 | 17.7 |
|  |  | CETCH<sup>†</sup> | 317 | 21 | 22.8 |
|  |  | rTCA<sup>‡</sup> | 267 | 25 | 19.1 |
|  |  | WLP<sup>‡</sup> | 262 | 25 | 18.8 |
|  | NADH | Calvin | 91.9 | 57 | 8.41 |
|  |  | rGlyP | 70.5 | 74 | 6.45 |
|  |  | CETCH<sup>†</sup> | 90.5 | 58 | 8.28 |
|  |  | rTCA<sup>‡</sup> | 76.2 | 68 | 6.97 |
|  |  | WLP<sup>‡</sup> | 74.8 | 70 | 6.84 |
| ClO<sub>4</sub><sup>-</sup> | QH<sub>2</sub> | Calvin | 194 | 34 | 13.9 |
|  |  | rGlyP | 154 | 44 | 11.0 |
|  |  | CETCH<sup>†</sup> | 191 | 35 | 13.8 |
|  |  | rTCA<sup>‡</sup> | 165 | 41 | 11.8 |
|  |  | WLP<sup>‡</sup> | 162 | 42 | 11.6 |
|  | NADH | Calvin | 83.1 | 63 | 7.60 |
|  |  | rGlyP | 66.0 | 79 | 6.04 |
|  |  | CETCH<sup>†</sup> | 82.0 | 64 | 7.50 |
|  |  | rTCA<sup>‡</sup> | 70.6 | 74 | 6.45 |
|  |  | WLP<sup>‡</sup> | 69.4 | 75 | 6.35 |
| SO<sub>4</sub><sup>2-</sup> | NADH | Calvin | 157 | 33 | 14.4 |
|  |  | rGlyP | 104 | 50 | 9.51 |
|  |  | CETCH<sup>†</sup> | 141 | 37 | 12.9 |
|  |  | rTCA<sup>‡</sup> | 118 | 44 | 10.8 |
|  |  | WLP<sup>‡</sup> | 115 | 45 | 10.5 |

<sup>†</sup>CETCH cycle has only been demonstrated *in vitro*
<sup>‡</sup>rTCA cycle and WLP require anoxic conditions
<sup>\*</sup>Values are derived from whole-number calculations so 3 significant figures are reported here arbitrarily

For a given electron sink and terminal electron acceptor (excepting O_2_ in the case of the rTCA cycle and WLP, which are oxygen-sensitive), the electron demand for the five CFPs we consider is ordered rGlyP<WLP<rTCA<CETCH<Calvin. Accordingly, the energy efficiency of carbon fixation is highest for the rGlyP, and the energy demand per unit of sucrose produced is lowest (Table 1). Using NADH as the electron sink results in a lower electron demand and higher energy efficiency than QH_2_. The former result is unsurprising since electrons deposited into the NADH pool are more energetic (have a lower redox potential) than those in the QH_2_ pool, and the latter indicates that reverse electron flow cannot overcome the efficiency challenge presented by starting with a low-energy electron donor. Using QH_2_ as the electron sink cannot support CO_2_-fixation with sulfate as the terminal electron acceptor because the sulfate redox couple (SO_4_^2-^/S^2-^, E^0^ = -0.22 V vs. SHE) is energetically uphill of the quinone pool (Q/QH_2_, E^0^ = -0.08 V vs. SHE). Of the four terminal electron acceptors we consider, O_2_ results in the smallest electron demand and the highest energy efficiency; the electron demand for each is ordered O_2_<ClO_4_^-^<NO_3_^-^<SO_4_^2-^. Although our analysis suggests that perchlorate would be the preferred electron acceptor over nitrate, nitrate is typically preferred by organisms that can respire each, possibly due to the risk of toxic intermediate buildup during the respiratory process.^53^

The highest energy efficiency (84%) and the lowest energy demand (5.67 kWh/kg) can be achieved by an organism that uses the NADH pool as a sink for cathode-derived electrons, O_2_ as a terminal electron acceptor, and the rGlyP to fix carbon (Table 1). However, metabolic energy efficiency alone is not enough to determine if a process is viable from a life cycle perspective. Upstream processes including electricity generation, CO_2_ capture, and electron acceptor (re-)generation all require explicit attention as drivers of life cycle impacts. Moreover, the productivity of the EMP system is extremely important since a lower productivity results in a high demand for electrolyzer materials per unit product over the process lifetime. We therefore developed a complete process model for dEMP processes (diagrammed in Fig. 1b) to understand full-system life cycle impacts, which we discuss next.

### Global warming potential calculations reveal target productivity

Four major factors contribute to the global warming potential (GWP) of sucrose production using dEMP (Fig. 1b): (1) direct air capture of CO_2_; (2) electron acceptor regeneration, which converts the reduced electron acceptor (*e.g*., N_2_) back into the oxidized form necessary for microbial respiration (*e.g*., NO_3_^-^); (3) electricity production, which supports direct air capture and electron acceptor regeneration processes in addition to driving the EMP reactor; and (4) reactor materials production, including the plastic reactor body, carbon paper electrodes supporting the anode metal catalyst and cathode biocatalyst, and the anode metal catalyst.

We calculated the GWP of these process components for each combination of electron sink, CFP, and terminal electron acceptor, as detailed in the supplementary information. We assume all energy is supplied by wind power, and we note that other forms of clean energy production (*e.g*., thin film photovoltaics) have roughly equal life cycle emissions and would therefore lead to similar results.^54^ In all cases, the GWP of direct air capture for the process is equivalent because we assume that all of the fixed carbon is diverted to sucrose production. The GWP of each terminal electron acceptor is different based on the specifics of the regeneration process, and is ordered O_2_ (0 kg CO_2,e_/kg O_2_) < SO_4_^2-^ (0.0056 kg CO_2,e_/kg Na_2_SO_4_) < ClO_4_^-^ (0.063 kg CO_2,e_/kg NaClO_4_^-^) < NO_3_^-^ (0.192 kg CO_2,e_/kg NaNO_3_). For the full process, the GWP contribution by the terminal electron acceptor also depends on the CFP, since less efficient CFPs (*e.g*., the Calvin cycle) require more ATP and NAD(P)H and therefore more of the terminal electron acceptor. Hence, microbes using the rGlyP not only consume less energy during the sucrose production process, but also consume less energy during terminal electron acceptor regeneration because they require a smaller amount of the terminal electron acceptor per unit sucrose.

The GWP contribution by reactor materials depends on the productivity of the EMP reactor because a higher productivity over the lifetime of the reactor results in more sucrose produced per unit of reactor materials. We therefore calculated the full-system GWP as a function of current density using the rGlyP to fix carbon and for each terminal electron acceptor and electron sink (Fig. 2). As the current density increases, the GWP decreases for each system considered until eventually approaching a plateau around 1000 mA/cm^2^ as the GWP contribution of reactor materials production becomes small with respect to that of energy demands to drive the reactor and terminal electron acceptor regeneration (Fig. 2).

**Figure 2.**
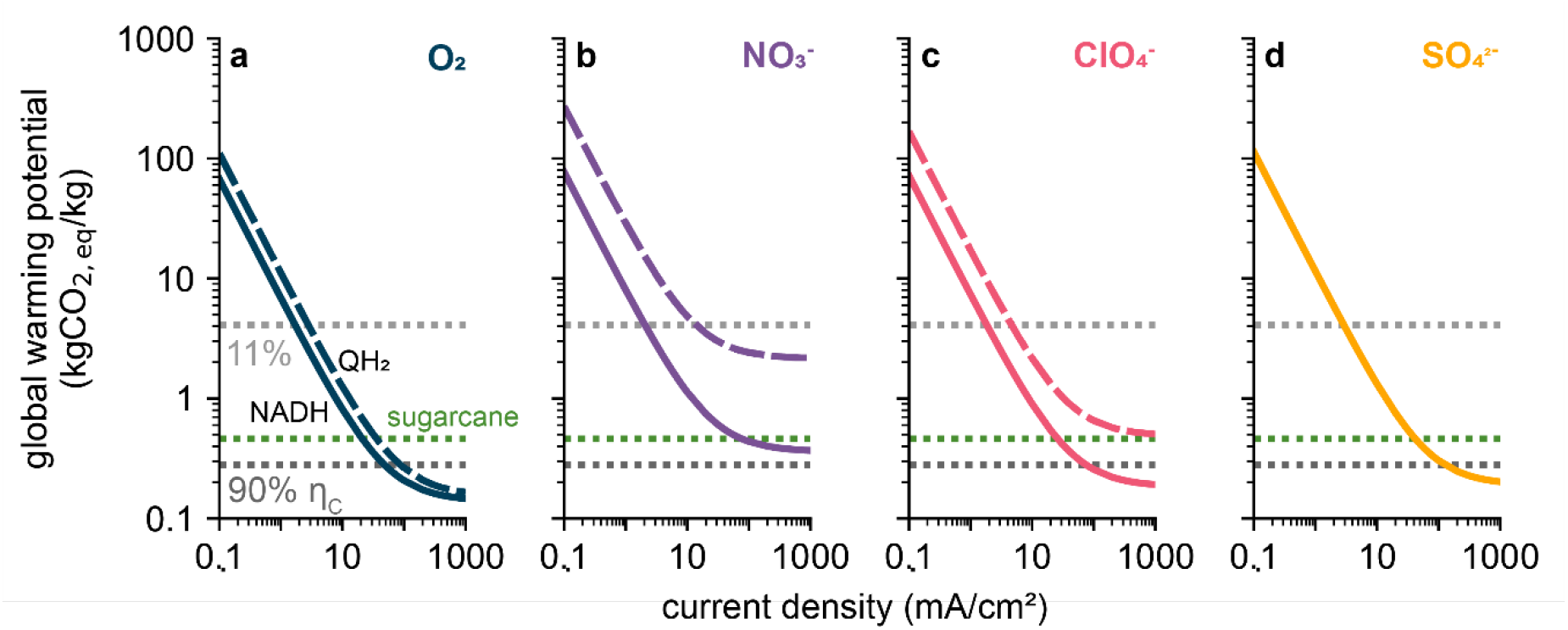
Global warming potential of direct electron transfer-based processes. Global warming potential of the production of sucrose as a function of current density for direct electron uptake using the reductive glycine pathway to fix carbon, the NADH (solid lines) or QH_2_ (dashed lines) pool as the electron sink, and (**a**) O_2_, (**b**) NO_3_^-^, (**c**) ClO_4_^-^, or (**d**) SO_4_^-^ as the terminal electron acceptor. Gray dotted lines in each panel correspond to the global warming potential of sucrose production for H_2_-mediated Knallgas bacteria operating at 11% (light gray) and 90% (dark gray) carbon efficiency. Green dotted line corresponds to the global warming potential of sucrose production from sugarcane. Panel (**d**) includes only electron uptake into the NADH pool (solid line) because the QH_2_ pool cannot support growth with SO_4_- as the terminal electron acceptor, as discussed in the main text.

Differences in the terminal electron acceptor drive large differences in the overall GWP: at 100 mA/cm^2^, the GWP of each process using NADH as the electron sink and the rGlyP to fix carbon is ordered (in units kg CO_2,e_/kg sucrose) O_2_ (0.21) < ClO_4_^-^ (0.26) < SO_4_^-^ (0.30) < NO_3_^-^ (0.44). Hence, although NO_3_^-^ enables more efficient carbon fixation than SO_4_^2-^(74% vs. 50%), the large GWP of nitrate regeneration causes this strategy to have a higher life cycle impact. Using QH_2_ as the electron sink instead of NADH further reinforces the importance of the terminal electron acceptor. Because deposition of electrons into the quinone pool requires reverse electron flow to generate NADH and ATP, significantly more terminal electron acceptor is required per unit sucrose (*e.g*., 1.12 kg NaNO_3_/kg sucrose for the NADH pool vs. 9.87 kg NaNO_3_/kg sucrose for the QH_2_ pool, using the rGlyP to fix carbon). Hence, the GWP of the full process is significantly higher if electrons are deposited into the quinone pool; at 100 mA/cm^2^, these values are (in units kg CO_2,e_/kg sucrose) 2.41 (vs. 0.44) for nitrate, and 0.65 (vs. 0.26) for perchlorate (Fig. 2). The difference between the QH_2_ and NADH pools as the electron sink is smaller when O_2_ is the terminal electron acceptor because O_2_ is regenerated naturally by the EMP process (via water oxidation at the anode), so the slight improvement (0.26 (QH_2_) vs. 0.21 (NADH)) is due to the increased efficiency of electron uptake and reduced electron demand through the NADH pool (Table 1).

Beyond considering differences between dEMP processes, we also wanted to benchmark this process strategy against alternative EMP options and traditional bioprocesses (*i.e*., those based on sugarcane). Our previous analysis demonstrated that H_2_-mediated EMP could be more effective (induce a lower GWP) than traditional bioprocesses for the production of microbial biomass (*e.g*. for use as single cell protein), industrial enzymes, and lactic acid.^5^ Here, we expanded that analysis to consider sucrose production from H_2_ with microbes using the Calvin cycle to fix CO_2_. We evaluated two different carbon efficiencies (fraction of fixed carbon diverted to sucrose): 11%, based on the reported value by Nangle *et al*.,^39^ and 90%, representing an optimistic, but feasible, upper bound set by cyanobacterial sucrose production.^45^ We also used sucrose production via sugarcane as an additional reference value, since this represents a dominant mode of sucrose production worldwide.

Our results demonstrate that H_2_-mediated EMP of sucrose can achieve a lower GWP than that of sugarcane-derived sucrose if the carbon efficiency is 90%. The lower carbon efficiency we considered, 11%, has a higher GWP than sugarcane-based production, although it would result in a reduced land occupation footprint.^39^ To break even (in terms of GWP) with the H_2_-mediated system, sucrose production based on direct electron uptake with O_2_ as the terminal electron acceptor, rGlyP as the carbon fixation pathway, and NADH as the electron sink would need to achieve a current density of ∼48.4 mA/cm^2^ (Fig. 2). With QH_2_ as the electron sink, the breakeven value is ∼88 mA/cm^2^ (Fig. 2, supplementary Fig. S1). Regardless of the electron sink, nitrate use as the terminal electron acceptor would prevent direct electron uptake-based systems from reaching parity with H_2_-mediated ones; this is due to the large GWP associated with nitrate regeneration. Although both perchlorate- and sulfate-reducing systems can achieve parity with H_2_-mediated sucrose production, this requires the use of NADH as the electron sink and current densities >76 mA/cm^2^ (ClO_4_^-^) and >128 mA/cm^2^ (SO_4_^2-^). Hence, this analysis suggests that microbes using the rGlyP to fix carbon, O_2_ as the terminal electron acceptor, and NADH as the electron sink represent the best option for dEMP, and that production based on this metabolic strategy may be able to outcompete alternative options.

However, this preliminary conclusion is troubled by three significant points of caution. First, we previously showed that standard dEMP systems are limited to <∼1.5 mA/cm^2^ by O_2_ diffusion through a fluid boundary layer and to <∼30 mA/cm^2^ by CO_2_ diffusion.^14^ Hence, architectures such as gas diffusion electrodes (GDEs), which enable an extremely high gas/liquid interfacial area,^55^ are necessary to achieve the high current density required for dEMP systems to compete with alternative options. Second, unavoidable efficiency losses within the EMP reactor, including kinetic overpotentials and ohmic losses in the electrolyte, will increase the energy demand per unit of sucrose produced, which will in turn increase the GWP of the overall process. Hence, a current density higher than 48 mA/cm^2^ is almost certainly necessary. Third, this high current density will cause significant pH and salinity increases in the reactor, so the microbial catalyst performing CO_2_-fixation must be able to not only withstand, but function optimally, in an extreme environment. We explore each of these considerations in detail in the following sections.

### Gas-phase transport and gas-liquid mass transfer in GDE architectures

Originally developed for fuel cells, GDEs have been successfully adopted for CO_2_ electroreduction.^56^ These systems can achieve current densities in excess of 300 mA/cm^2^ for the abiotic reduction of CO_2_ to formate, carbon monoxide, and ethylene, and have also been employed to enhance the gas-liquid mass transfer rate in microbial electrosynthesis reactors.^56–58^ Comprehensive physical modeling of GDEs and their application in membrane electrode assemblies have been carried out for abiotic catalysts,^55,59,60^ demonstrating that CO_2_ gas-liquid mass transfer does not limit the productivity of these systems. However, in abiotic GDE architectures, the catalyst layer is typically only ∼5 μm thick.^55,59^ In contrast, for a bio-GDE with a microbial volume fraction of 0.52, a rate-limiting enzymatic turnover of 100 s^-1^, and an electron demand of 62 mol e^-^/mol sucrose (corresponding to carbon fixation through the rGlyP with formate dehydrogenase as the rate-limiting enzyme,^61^ electron uptake into the NADH pool, and O_2_ reduction as the terminal electron acceptor), the biocatalyst layer (bCL) would need to be >170 μm thick to enable a current density >100 mA/cm^2^. This >30-fold difference in catalyst layer thickness could induce significant differences in the transport behavior of bio-GDE systems. We therefore wanted to confirm that gas phase transport and gas liquid mass transfer would not limit the current density of a dEMP system (diagrammed in Fig. 1c).

We considered two relevant cases for this system: one with an extremely high surface area (1 × 10^6^ m^2^/m^3^) with a monolayer of cells distributed evenly along the carbon support, and one with a much lower surface area (5.6 × 10^5^ m^2^/m^3^) and a substantially thicker biofilm (10 μm). The former case approximates the biotic analogue of GDEs designed for CO_2_ electrolysis with a thin catalyst layer and ultra-high specific surface area, while the latter is similar to the architecture proposed previously for biological systems.^16^

For both bio-GDE architectures, the liquid-phase concentration of CO_2_ and O_2_ retain >98% of their initial values throughout the length of a 325 μm-thick biocatalyst layer operating at its limiting current density (Fig. S2). Both the total pressure and individual partial pressure decrease by <0.1% for each architecture (data not shown). Hence, neither gas phase transport nor gas-liquid mass transfer limit the achievable current density in bio-GDEs. We continue our analysis here with the first case (high surface area, monolayer of cells) because the high surface area support enables a higher biomass density throughout the electrode, reducing the total bio-GDE thickness necessary to achieve a given productivity. In addition, restricting biofilm formation to a monolayer of cells minimizes the electron transport distance and relaxes demands on the conductivity of the extracellular matrix or conductive membrane extrusions or pili, which together may enable a higher fraction of cells to be metabolically active.^62^

### Chemical species transport, pH, and salinity

Because gas-liquid mass transfer limitations can be overcome by employing a bio-GDE architecture, we continued our analysis by developing a comprehensive bio-electrochemical model of the bio-GDE EMP system. Since carbon fixation with the rGlyP coupled to aerobic respiration can achieve the highest efficiency and lowest GWP (Table 1, Fig. 2), our analysis focuses on this system.

We used our model, which describes mass transport, (bio)electrochemical and acid-base thermodynamics and kinetics, and gas-liquid mass transfer (see Supplementary Information for complete modeling details), to calculate the total system voltage necessary to achieve a given current density. In addition, because CO_2_ fixation results in the net consumption of protons, the pH throughout the biocatalyst layer (bCL) will increase as the current density increases. An increased pH will promote the formation of bicarbonate and carbonate species throughout the bCL, both of which are negatively charged, so positive ions (we used Na^+^ in our model) will be drawn into the liquid phase in the bCL to maintain charge neutrality. The net effect will be a substantial increase in the salinity throughout the bCL. Because microbes are sensitive to both pH and salinity extremes, we calculated the average pH and salinity developed throughout the bCL during steady-state operation.

When NADH is used as the electron sink, the total system voltage necessary to drive the reaction (which includes the thermodynamic potential, kinetic overpotentials, and Nernst and ohmic losses) increases from <1.8 V to ∼2.8 V as the current density is increased from <5 mA/cm^2^ to 150 mA/cm^2^ (Fig. 3a). The total voltage is slightly lower when electrons are deposited into the QH_2_ pool; this is mostly due to the difference between the thermodynamic potential necessary for quinone reduction and that for NAD^+^ reduction (Fig. 3a). However, an equal current density for the two systems does not result in an equal sucrose productivity because more electrons are required per sucrose when electrons are deposited into the quinone pool (Table 1).

**Figure 3.**
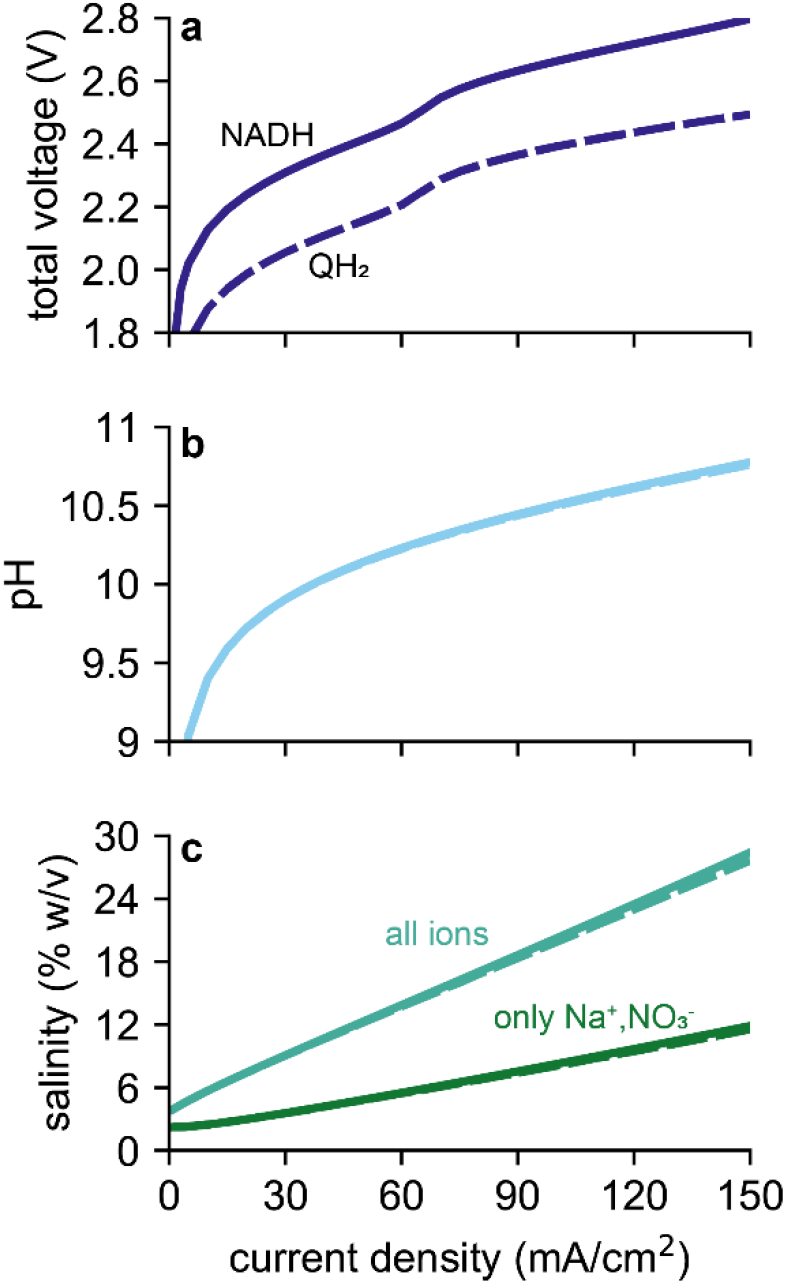
Reactor conditions. (**a**) total system voltage, (**b**) average pH in the biocatalyst layer (bCL), and (**c**) average salinity (% w/v) in the bCL as a function of current density for bacteria fixing carbon with the reductive glycine pathway, using O_2_ as the terminal electron acceptor, and depositing cathode-supplied electrons into the NADH (solid lines) or QH_2_ (dashed lines) pool.

Both the average pH and average salinity increase, as expected, as the current density increases (Fig. 3b, c). At 48 mA/cm^2^, the average pH is 10.1, with an average total salinity of 11.97% (w/v). At 150 mA/cm^2^, these values increase to 10.8 and 28.4%, respectively (Fig. 3b, c). Hence, in the current density regime that is necessary for ecologically viable production (*i.e*., a GWP near the GWP of H_2_-mediated sucrose production), the microbe would need to withstand high pHs and salinities. Because well-characterized electroautotrophs (*e.g*., *R. palustris*) are neutrophilic and non-halophilic,^25^ tolerance to extreme conditions would need to be engineered into these organisms. Alternately, known extremophiles could be tested for electroautotrophic capability or engineered to contain the necessary metabolic machinery for electroautotrophy.

A limitation to our model is that we have used dilute-solution theory to describe the transport of liquid-phase species. This assumption may break down at high current densities as concentrations of carbonate and sodium species increase to well above 1 M.^55,63^ More general concentrated-solution theory would account for additional drag experienced by diffusing species arising from their interactions with other species and would correct the thermodynamic driving force for diffusion and migration to account for these terms.^63^ However, the general trends we report here are not expected to change significantly with corrected parameters, and would likely operate to increase the voltage, pH, and salinity at a given current density by no greater than ∼10%. We also note that other models of CO_2_ electroreduction have used dilute-solution theory with reasonable accuracy at significantly higher species concentrations than what we observe here.^55^

### Global warming potential for a highly-engineered system

Assuming that an organism with the pH- and salinity-tolerance necessary to achieve high current density can be identified or engineered, we calculated the GWP associated with sucrose production using our full electrochemical system model to predict the energy demand per unit of sucrose as a function of the current density (Fig. 4a). When reactor inefficiencies (due, for example, to kinetic or Ohmic overpotentials) are included, the current density necessary to match H_2_-mediated sucrose production (at 90% carbon efficiency) increases from ∼48 mA/cm^2^ (Fig. 2) to 91 mA/cm^2^ (Fig. 4a). Interestingly, the bio-GDE system using QH_2_ as the electron sink cannot reach parity with the H_2_-mediated case because of the higher electron demand (Fig. 4a).

**Figure 4.**
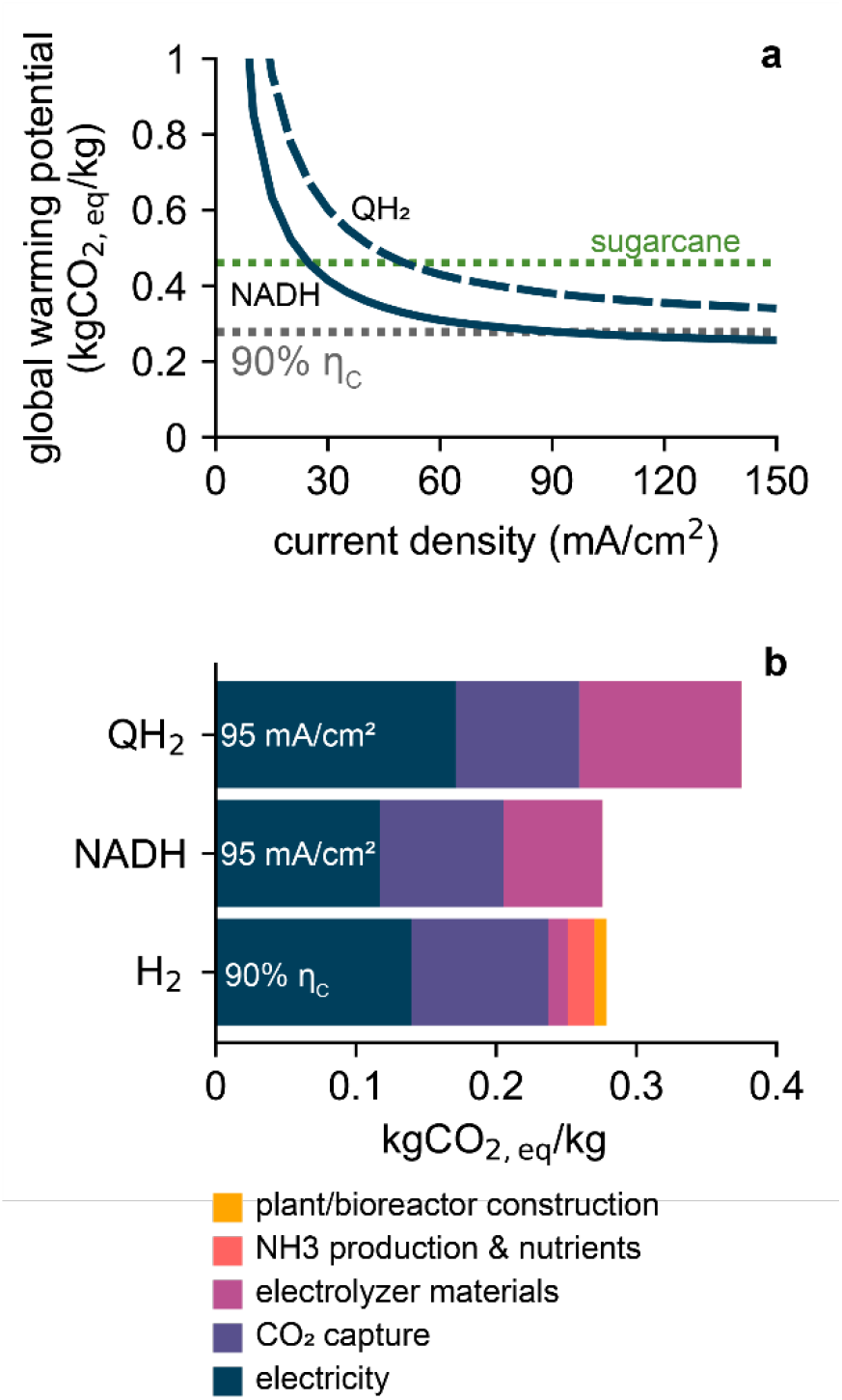
Modeled global warming potential of direct electron uptake-based processes. (**a**) Global warming potential of sucrose production for direct electron uptake as a function of current density for bacteria using the reductive glycine pathway to fix carbon, the NADH (solid lines) or QH_2_ (dashed lines) as an electron sink and O_2_ as the terminal electron acceptor. Gray dotted line corresponds to the global warming potential of the H_2_-mediated Knallgas system operating at 90% carbon efficiency. Green dotted line corresponds to the global warming potential of sucrose production from sugarcane. (**b**) Global warming potential for the three EMP systems considered in (**a**) broken down by process category.

We break down the GWP for each EMP process by subprocess category in Fig. 4b. The GWP associated with electrolyzer materials for direct electron uptake-based systems is significantly higher than that for H_2_-mediated production (Fig. 4b); this is because dEMP systems operate at a significantly lower current density (here, 95 mA/cm^2^ *vs*. 1000 mA/cm^2^). Although the GWP associated with electricity demand for direct systems is lower when the NADH pool is used as the electron sink (because we’ve assumed 100% of fixed CO_2_ is diverted to sucrose), the electricity demand when QH_2_ is used as the electron sink is substantially higher (Fig. 4b). We attribute this to the higher electron demand per unit sucrose (Table 1). The higher electron demand also causes the GWP associated with reactor (electrolyzer) materials production to be higher at equivalent current densities because the productivity is lower (Fig. 4b). These twin effects prevent QH_2_ use as the electron sink from reaching parity with the H_2_-mediated system. At current densities high enough to minimize GWP contribution from electrolyzer materials, the GWP associated with the power demand necessary to achieve that current density (along with the GWP from CO_2_ capture) is already higher than the full-system GWP of H_2_-mediated production. In addition to this practical limitation, electron uptake into the QH_2_ pool has a higher energy demand (for the Calvin cycle, 8.70 kWh/kg) than H_2_-mediated carbon fixation (∼7.8 kWh/kg) for sucrose production when each process operates at its thermodynamic maximum efficiency (Table 1). Hence, only an organism that uses NADH as the electron sink can outcompete H_2_-mediated production, and even then, only marginally. This presents a significant challenge to dEMP systems since electron conduits that interface with the NADH pool are uncharacterized to date.

### Charting the narrow course for direct electron uptake-based EMP

Throughout the analysis in this work, we have made a series of optimistic assumptions about the feasibility of dEMP processes. First, we assumed that an organism with the necessary metabolic components (a high-efficiency and high-rate carbon fixation pathway, an electron conduit that interfaces with the NADH pool, and aerobic respiratory capacity) could be identified or engineered wherein each physiological module functions at its maximal efficiency. Second, we assumed this organism could be effectively attached to a high-surface area gas diffusion electrode in a dense monolayer, and that this bio-GDE could be engineered to have optimal wetting characteristics to maximize liquid and gas transport. Third, we assumed that the microbe would not need to divert any fixed carbon to maintain cellular activity (*i.e*., that it could achieve 100% carbon efficiency to the desired product, sucrose, and that it would not need any additional elemental resources including nitrogen or phosphorous). Fourth, we assumed that this organism could function optimally at pH (>10.4) and salinity (∼18.8%, composed of ∼3.2 M Na^+^ and ∼1.87 M dissolved inorganic carbon) extremes with no loss in efficiency or productivity.

Could an organism with these characteristics be engineered? Existing or predicted electroautotrophs function optimally in near-neutral pH ranges and low salt concentrations. Although adaptive laboratory evolution strategies have enabled neutrophilic and/or non-halophilic organisms to grow at moderately alkaline or saline conditions, these efforts have so far only enabled tolerance up to pH <9.5 and <1 M Na^+^.^64,65^ Engineering further tolerance to extreme conditions is probably challenging since most cellular machinery would likely have to adapt to increased cytoplasmic pH and osmolarity.

A more effective solution could be to engineer an existing haloalkaliphile to use electrons accepted from a cathode as an energy source for autotrophic growth. Organisms requiring the fewest modifications to their central metabolism would be best-suited to this task. Autotrophic, aerobic haloalkaliphiles would require only the introduction of an electron conduit, a feat that has previously been accomplished in *E. coli*.^66^ Among cyanobacteria, *Euhalothece* sp. ZM001 grows optimally at a pH of ∼10, but requires ∼0.5 M Na^+^ and <1.5 M dissolved inorganic carbon.^67^ Among chemolithotrophs, the alphaproteobacterial strain AHO 1 (closely related to *Roseinatronobacter thiooxidans*) grows optimally in the pH range 9.5 – 9.8 and continues H_2_ oxidation up to pH 11.^68^ However, the salt tolerance of this organism is <1 M Na^+^, and other chemolithotrophs with higher salt tolerances are obligately anaerobic.^69^ Hence, few promising cultured candidates exist based on these criteria, so expanded efforts to culture autotrophs with these desired phenotypes would be required in parallel to engineering efforts.

Moving away from known autotrophs, several extreme haloalkaliphilic chemoorganotrophs across both bacterial and archaeal domains exist that respire O_2_ and grow optimally in extremely high pH and salinity regimes. Within bacteria, *Halomonas* and *Salinicoccus* genera contain obligately aerobic halophiles that can withstand Na^+^ concentrations in excess of 4.2 M and pH in excess of 11. However, optimal salinities (*e.g*., ∼1.5 M Na^+^ for *Halomonas cupida*) are well below the predicted conditions in the bio-GDE.^70^ Within archaea, both *Halorubrum* and *Natronolimnobius* genera contain obligately aerobic halophiles that can withstand Na^+^ concentrations up to 5.2 M and pH up to 11 while maintaining growth.^71^ In these species, the optimal growth salinity (*e.g*., 4.1 M Na^+^ for *Halorubrum alkaliphilum*)^72^ are above what is predicted in the bio-GDE, suggesting that they would be good microbial chassis candidates. None of these species, however, are known to fix CO_2_ nor express characterized electron conduits,^14^ although we note that archaeal electron conduits are under-characterized as compared to those of bacterial origin. This indicates that both a carbon fixation pathway (rGlyP) and an electron conduit (that interfaces with the NADH pool) would be necessary to introduce into one of these organisms. Although, in principle, this is possible, as carbon fixation pathways and an electron conduit have been functionalized in *E. coli*,^66,73,74^ to our knowledge, no genetic tools for any of these extremophilic organisms currently exist. Moreover, a complete electron conduit (*e.g*., from *Methanosarcina barkeri*)^21^ that interfaces with the NADH pool would need to be elaborated prior to its functional reconstitution in a different host organism.

Could improved reactor design ease the requirements on the microbial chassis? Our previous analysis demonstrated that standard reactor designs (Fig. 5) severely restricted the achievable current density in direct electron transfer-based EMP systems.^14^ Here, we have shown that deploying a bio-GDE architecture can overcome this limitation. However, at the high current density necessary to break even with H_2_-mediated systems, microbes experience pH and salinity extremes that make this strategy extremely challenging (Fig. 3, 4).

**Figure 5.**
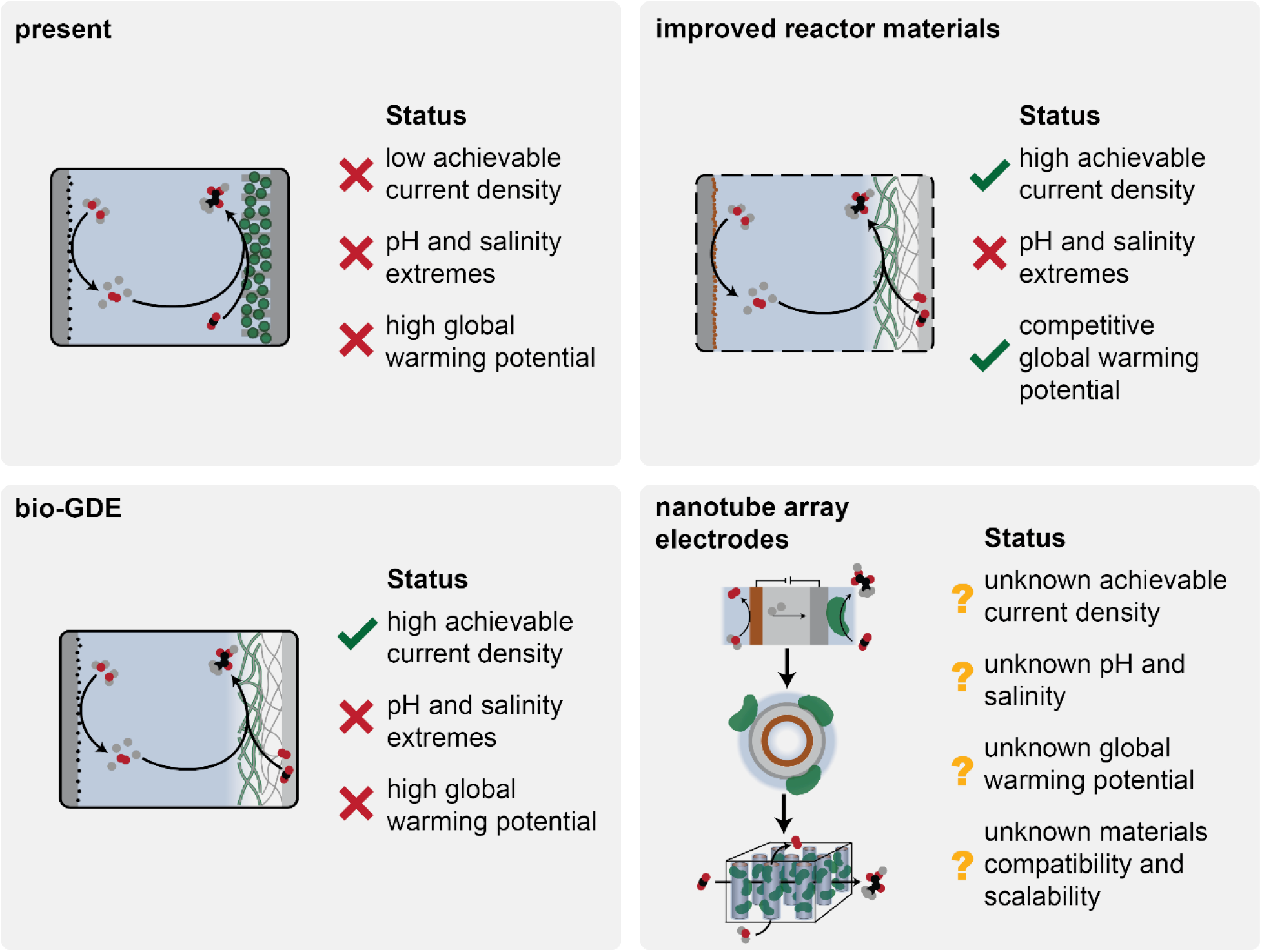
Potential reactor designs for improved performance. Current EMP reactors rely on bulk gas-liquid mass transfer, severely restricting the achievable current density. Deploying gas diffusion electrode (GDE) architectures overcome this limitation, but less resource-intensive materials are necessary, and microbes will experience pH and salinity extremes. Novel nanotube array electrodes^75,76^ have been proposed to overcome these limitations, but neither their potential nor practicality are understood.

Modifying electrode materials to be less resource intensive may partially address this issue (Fig. 5). In our analysis, we have assumed that electrolyzers are constructed from *poly*-methyl methacrylate (PMMA), a common electrolyzer body material, and we used an IrO_2_/SnO_2_ anode following our previous report on mediated EMP systems.^5^ PMMA comprises ∼70% of the GWP of the electrolyzer materials, and its impact could be reduced by a factor of ∼3 if it was replaced with a renewable-sourced plastic such as *poly*-lactic acid.^5^ This would reduce the breakeven current density (when electrons are transferred to the NADH pool) to ∼50 mA/cm^2^, corresponding to an average pH of ∼10.1 with ∼2 M Na^+^ and ∼1.2 M dissolved inorganic carbon in the bio-GDE (Fig. 3). Further replacing the IrO_2_/SnO_2_ anode with earth-abundant catalysts such as NiFe hydroxides^75^ would decrease the breakeven current density to ∼25-30 mA/cm^2^, corresponding to a pH of ∼9.9 with ∼1.3 M Na^+^ and ∼0.9 M dissolved inorganic carbon (Fig. 3). We note, however, that earth-abundant catalysts would likely result in a higher overpotential associated with the water oxidation reaction. Some haloalkaliphilic autotrophs (*e.g*., *Euhalothece* sp. ZM001) may be adaptable to these conditions;^67^ but the native Calvin cycle would need to be replaced with the reductive glycine pathway^76^ and an NADH pool-interfacing electron conduit would need to be functionally expressed, in addition to further metabolic engineering necessary to shunt carbon towards the desired product. Notably, regardless of improvements to reactor materials, electron conduits that interface with the QH_2_ pool still cannot support a dEMP system that breaks-even with the H_2_-mediated system.

Novel electrode designs relying on nanotube arrays have recently been proposed for EMP applications (Fig. 5).^77,78^ In this design, the nanotube comprises the complete electrolyzer “stack”, reducing the proton transport distance to tens of nanometers and potentially enabling convective transport of chemical reactants to the anode and biocathode surfaces (Fig. 5).^77,78^ This approach could limit the pH gradient and salt buildup in the vicinity of the microbes while maintaining a high achievable current density. However, several design criteria must be met simultaneously in this system. First, the nanotube inner diameter and spacing must enable low pressure-drop transport of reacting species through the array. Second, the nanoscale membrane must conduct protons rapidly while blocking electron transport/tunneling between electrodes during operation at 1.5–2.5 V potential differences (Fig. 3a). Third, the array must enable robust microbial attachment even in the presence of significant local fluid velocities that could cause detachment. Fourth, nanowires are commonly deployed as antimicrobial agents,^79,80^ so structure-function relationships must be determined to ensure microbe viability. Fifth, wetting characteristics must be optimized such that microbes can survive in a moist environment but without such a thick liquid film that gas exchange is inhibited and the achievable current density is limited. Finally, the nanotube arrays must be reliably synthesized at scale, and array fabrication must minimize resource intensity relative to much more easily constructed water electrolyzers. Currently, it is unclear which, if any, of these requirements are practical.

Given the significant challenges facing the design and engineering of any dEMP system, it is worth revisiting the central motivations driving the development of such systems. dEMP systems promise to achieve higher efficiency and lower resource intensity than mediated systems. Higher efficiency is predicted because NADH has a less negative redox potential than H_2_ (−320 mV *vs*. -411 mV), reducing the minimum thermodynamic energy demand by ∼7–8% when H_2_O/O_2_ (+818 mV) is the redox couple. Moreover, avoiding a mediator molecule eliminates potential losses associated with imperfect utilization of the mediator. Lower resource intensity is assumed because integrating multiple functionalities into a single reactor should reduce the overall balance-of-systems intensity. However, our analysis indicates that the resource intensity of the direct electron transfer-based EMP system is higher than that of the H_2_- mediated system due to the lower achievable current density (Fig. 4). The marginal potential benefit in energy efficiency, then, must act to offset the higher resource intensity.

Our analysis indicates that developing practical dEMP processes would require overcoming substantial hurdles in microbial discovery and engineering, (nano)materials synthesis, and reactor design and scale-up. In contrast, H_2_-mediated systems rely on industrially-established water electrolysis^81^ and gas fermentation technologies^82^ that readily operate at scale with high efficiency. We suggest, therefore, that research and development towards practical applications of dEMP should be benchmarked against mediated EMP processes. This analysis, however, should not discourage efforts to study the physiology of extracellular electron transfer in microbes in general. Electron transfer mechanisms appear to be widespread in nature and of significant ecological importance.^10^ Moreover, applications in toxicant biosensing,^83^ wastewater treatment,^84^ and environmental remediation^85^ are highly promising, but these efforts are subject to several issues beyond the technological.^86^

## Author Contributions

**Conceptualization:** A.J.A., J.D.A., J.M.H.; **Data curation:** A.J.A., J.D.A.; **Formal analysis:** A.J.A.; **Funding acquisition:** A.P.A. **Methodology:** A.J.A., J.D.A., J.M.H.; **Software:** A.J.A.; **Supervision:** A.P.A.; **Visualization:** A.J.A.; **Writing – original draft:** A.J.A.; **Writing – review & editing:** A.J.A., J.D.A., J.M.H, A.P.A.

## Acknowledgements

This work was supported by the Center for the Utilization of Biological Engineering in Space (CUBES, https://cubes.space), a NASA Space Technology Research Institute (grant number NNX17AJ31G). A.J.A. is supported by an NSF Graduate Research Fellowship under grant number DGE 1752814. We thank Prof. Caroline Ajo-Franklin (Rice University) and Dr. Heinz Frei (Lawrence Berkeley National Laboratory) for useful discussions on direct electron transfer systems, Helen Bergstrom (UC Berkeley) for useful discussions on electrochemistry, Marisa Watanabe for advice on schematics, and Dr. Paul Tol (Netherlands Institute for Space Research) for a helpful reference on accessible color schemes (https://personal.sron.nl/~pault/). We acknowledge this research was performed on unceded land of the Chochenyo-speaking Ohlone people.

## Notes

### Competing Interest Statement

The authors have declared no competing interest.

