## Supplementary Information for "Charting a narrow course for direct electron uptake-facilitated electromicrobial production"

#### Supplementary Figures

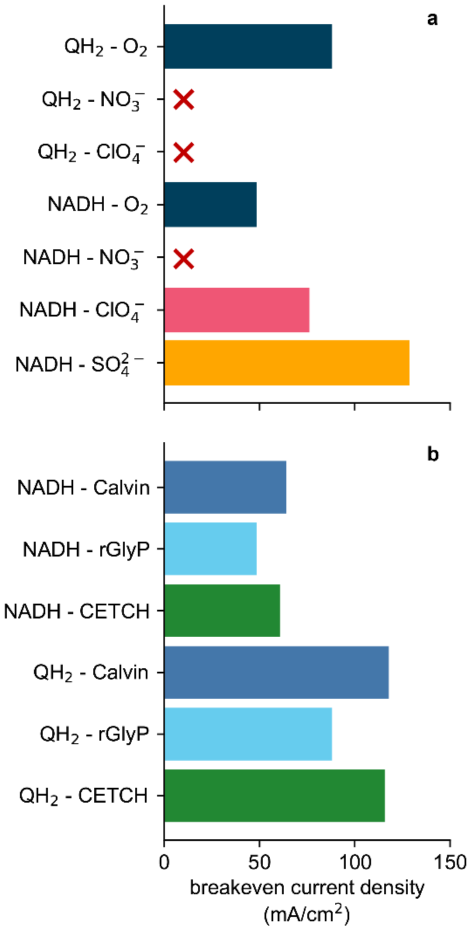

**Figure S1. Breakeven current densities.** Current density at which the GWP of DET-based EMP is equal to the GWP of $H_2$-mediated EMP operating at 90% carbon efficiency. Red "x"s indicate that a given DET metabolic strategy is unable to achieve parity with $H_2$-mediated system. All bars in (**a**) assume the reductive glycine pathway is used to fix carbon. All bars in (**b**) assume $O_2$ is the terminal electron acceptor. Note that these values do not consider the additional energy demand necessary to overcome efficiency losses in the DET reactor.

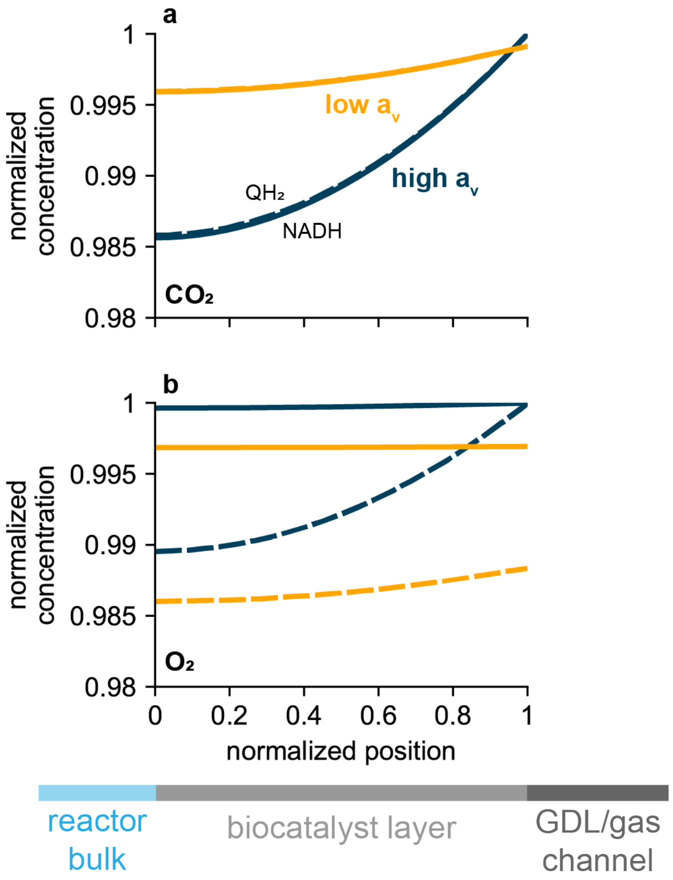

**Figure S2. $CO_2$ and $O_2$ transport in the bio-GDE.** Liquid phase (**a**) $CO_2$ and (**b**) $O_2$ concentration as a function of normalized position within the biocatalyst layer for the low specific surface area (yellow curves) and high specific surface area (blue curves) GDE cases. Microbes are assumed to be fixing carbon using the reductive glycine pathway, with $O_2$ as the terminal electron acceptor and $QH_2$ (dashed curves) or NADH (solid curves) as the electron sink. "High" and "low" specific surface areas correspond to $1 \times 10^6$ $m^{-1}$ and $5.6 \times 10^4$ $m^{-1}$, as described in the main text. These calculations use a 325 μm biocatalyst layer; the projected-area current densities are 173 mA/cm$^2$ (NADH, high $a_v$), 283 mA/cm$^2$ ($QH_2$, high $a_v$), 97 mA/cm$^2$ (NADH, low $a_v$), and 158 mA/cm$^2$ ($QH_2$, low $a_v$).

### Methods

**Stoichiometry and energetic analysis of electroautotrophic metabolisms**

**Carbon fixation pathways**

Different carbon fixation pathways (CFPs) generate different molecules as the primary product; we normalized each CFP to pyruvate as a common molecular intermediate, resulting in

Calvin cycle:[1,2]

$$3CO_2 + 5NADH + 7ATP + 4H^+ \rightarrow C_3H_3O_3^- + 5NAD^+ + 7(ADP + P_i) + 3H_2O \quad (1)$$

Reductive tricarboxylic acid (rTCA) cycle:[1,2]

$$\begin{aligned} &3CO_2 + 3NADH + 2ATP + 2Fd^{2-} + 6H^+ \\ &\quad \rightarrow C_3H_3O_3^- + 3NAD^+ + 2(ADP + P_i) + 2Fd + 3H_2O \end{aligned} \quad (2)$$

Reductive glycine pathway (rGlyP):[3,4]

$$3CO_2 + 5NADH + 2ATP + 4H^+ \rightarrow C_3H_3O_3^- + 5NAD^+ + 2(ADP + P_i) + 3H_2O \quad (3)$$

Crotonyl-CoA/ethylmalonyl-CoA/hydroxybutyryl-CoA (CETCH) cycle:[5]

$$\begin{aligned} &3CO_2 + 7NADH + 4ATP + 2FAD + 6H^+ \\ &\quad \rightarrow C_3H_3O_3^- + 7NAD^+ + 2FADH_2 + 4(ADP + P_i) + 3H_2O \end{aligned} \quad (4)$$

Wood-Ljungdahl pathway (WLP):[1,2]

$$\begin{aligned} &3CO_2 + 2NADH + 1ATP + 3Fd^{2-} + 7H^+ \\ &\quad \rightarrow C_3H_3O_3^- + 2NAD^+ + 1(ADP + P_i) + 3Fd + 3H_2O \end{aligned} \quad (5)$$

**Sucrose production**

Sucrose is generated by the conversion of primary products from CFPs via gluconeogenesis. Using pyruvate as a basis,

$$\begin{aligned} &4C_3H_3O_3^- + 4NADH + 13ATP + 8H^+ \\ &\quad \rightarrow C_{12}H_{22}O_{11} + 4NAD^+ + 13(ADP + P_i) + 1H_2O \end{aligned} \quad (6)$$

Note that here we have assumed ATP, GTP, and UTP are interchangeable.

**Direct electron uptake**

We consider two potential mechanisms by which electrons are transferred from the cathode into energy carrier pools in the cell. In the first, based on the MtrCAB/CctA/CymA conduit from *Shewanella oneidensis*,[6–8] electrons are transferred into the quinone pool according to

$$Q + 2H^+_{in} + 2e^- \rightarrow QH_2 \quad (7)$$

Most characterized mechanisms for electron uptake rely on the quinone pool as an electron sink.[9,10] Still, some sulfate-reducing microbes (SRMs),[11] methanogens,[12,13] and acetogens[14] may be able to grow on electrons directly without $H_2$ (or an equivalent) as a mediator. This metabolism would not be thermodynamically possible if electrons were deposited into the quinone pool: quinone oxidation cannot drive proton motive force generation with sulfate as the terminal electron acceptor and energy conservation in acetogens and methanogens requires direct NADH generation. Therefore, these bacteria must rely on some mechanism for electron uptake that follows the net reaction

$$NAD^+ + H^+_{in} + 2e^- \rightarrow NADH \quad (8)$$

These reactions require electrons at different potentials. For electron uptake into the quinone pool, we assume electrons are supplied at -100 mV *vs.* SHE because this is roughly the midpoint redox potential of MtrC[15] and is sufficiently electropositive to reduce the quinone pool (midpoint redox potential of >-80 mV *vs.* SHE). Following a similar strategy, we use -350 mV *vs.* SHE for electrons deposited into the $NAD^+$/NADH pool because the midpoint redox potential is ~-320 mV *vs.* SHE, and because -350 mV represents a rough average of the range of putative electron uptake mechanisms linked to SRMs and methanogens.[11,12]

**Energy carrier regeneration**

When electrons are deposited into the quinone pool, NADH is (re)generated by reverse operation of complex I in the electron transport chain (ETC):

$$NAD^+ + 4H^+_{out} + QH_2 \rightarrow NADH + 5H^+_{in} + Q \quad (9)$$

ATP is (re)generated by the action of ATP synthase according to

$$ADP + P_i + 3H^+_{out} \rightarrow ATP + 3H^+_{in} \quad (10)$$

We use the stoichiometry of the Rnf complex in, for example, *Clostridium ljungdahlii* to calculate ferredoxin regeneration:[16,17]

$$Fd + NADH + 2H^+_{out} \rightarrow Fd^{2-} + NAD^+ + 3H^+_{in} \quad (11)$$

**Respiration**

*$O_2$ as the terminal electron acceptor*

In the case where electrons taken up from the cathode are deposited into the quinone pool, complex III in the ETC releases 4 protons into the periplasmic space and liberates two electrons for later $O_2$ reduction according to

$$QH_2 + 2H^+_{in} \rightarrow Q + 4H^+_{out} + 2e^- \quad (12)$$

The two liberated electrons are transported by *c*-type cytochromes to ETC complex IV, transporting two additional protons across the inner membrane:

$$\frac{1}{2}O_2 + 4H^+_{in} + 2e^- \rightarrow H_2O + 2H^+_{out} \quad (13)$$

Hence, the overall reaction for quinol oxidation coupled to $O_2$ reduction is written as

$$QH_2 + \frac{1}{2}O_2 + 6H^+_{in} \rightarrow Q + H_2O + 6H^+_{out} \quad (14)$$

When electrons are deposited into the NADH pool, we use the standard description of oxidative phosphorylation assuming a $P/O$ ratio of 3 (*i.e.,* assuming the best-possible energy conservation):

$$NADH + H^+_{in} + \frac{1}{2}O_2 + 3ADP + 3P_i \rightarrow NAD^+ + H_2O + 3ATP \quad (15)$$

*$NO_3^-$ as the terminal electron acceptor*

Nitrate is reduced first to nitrite by quinol oxidation via the NarGHI complex:[18,19]

$$NO_3^- + 2H^+_{in} + QH_2 \rightarrow NO_2^- + H_2O + 2H^+_{out} + Q \quad (16)$$

Quinol is also oxidized to liberate electrons necessary for reducing nitrite all the way to $N_2$ following

$$\begin{gathered} 1.5QH_2 + 3H^+_{in} \rightarrow 1.5Q + 6H^+_{out} + 3e^- \\ NO_2^- + 2H^+_{out} + e^- \rightarrow NO + H_2O \end{gathered} \quad (17)$$

$$NO + H^+_{out} + e^- \rightarrow \frac{1}{2}N_2O + \frac{1}{2}H_2O$$

$$\frac{1}{2}N_2O + H^+_{out} + e^- \rightarrow \frac{1}{2}N_2 + \frac{1}{2}H_2O$$

Partial denitrification (*e.g.*, to nitrite) is also possible, but we neglect this possibility for two reasons related to the resulting build-up of the denitrification product. First, high nitrite concentration is typically toxic to cells.[20] Second, build-up of nitrite or other soluble terminal electron acceptors can negatively affect the thermodynamics of the terminal electron acceptor process. Hence, the overall reaction for quinol oxidation coupled to complete denitrification from nitrate is given by

$$2.5QH_2 + NO_3^- + 5H^+_{in} \rightarrow 2.5Q + 0.5N_2 + 3H_2O + 4H^+_{out} \qquad (18)$$

When electrons taken up from the cathode are deposited into the $NAD^+$/NADH pool, complex I in the ETC transfers electrons from NADH into the quinone pool, written as

$$NADH + 5H^+_{in} + Q \rightarrow NAD^+ + 4H^+_{out} + QH_2 \qquad (19)$$

Hence, the overall proton motive force-generating reaction associated with $NO_3^-$ reduction in this case is given by

$$2.5NADH + NO_3^- + 17.5H^+_{in} \rightarrow 2.5\ NAD^+ + 0.5N_2 + 3H_2O + 14H^+_{out} \qquad (20)$$

*$ClO_4^-$ as the terminal electron acceptor*

Perchlorate is first reduced to chlorite by the action of PcrQ/O/C/B/A according to[21,22]

$$2QH_2 + ClO_4^- \rightarrow 2Q + 2H_2O + ClO_2^- \qquad (21)$$

Chlorite dismutase splits $ClO_2^-$ into $Cl^-$ and $O_2$ without energy conservation. The $O_2$ is then reduced by complex IV of the ETC,

$$O_2 + 8H^+_{in} + 4e^- \rightarrow 2H_2O + 4H^+_{out} \qquad (22)$$

using electrons liberated by the oxidation of two extra quinols in complex III of the ETC:

$$2QH_2 + 4H^+_{in} \rightarrow 2Q + 8H^+_{out} + 4e^- \qquad (23)$$

Hence, the overall reaction for quinol oxidation coupled to perchlorate reduction is given by

$$4QH_2 + ClO_4^- + 12H^+_{in} \rightarrow 4Q + Cl^- + 4H_2O + 12H^+_{out} \qquad (24)$$

When electrons from the cathode are deposited into the $NAD^+$/NADH pool, complex I in the ETC transfers electrons from NADH into the quinone pool, written as

$$NADH + 5H^+_{in} + Q \rightarrow NAD^+ + 4H^+_{out} + QH_2 \qquad (25)$$

The remainder of the respiratory system is the same as written above, resulting in an overall reaction given by

$$4NADH + ClO_4^- + 32H_{in}^+ \rightarrow 4NAD^+ + Cl^- + 4H_2O + 28H_{out}^+ \qquad (26)$$

*$SO_4^-$ as the terminal electron acceptor*
Electrons taken up from a cathode that are deposited into the quinone pool cannot be used to generate a proton motive force when quinol oxidation is coupled to sulfate reduction because the quinone pool (~-80 mV *vs.* SHE) is more electronegative than the $SO_4^{2-}/H_2S$ redox couple (~-220 mV *vs.* SHE). Hence, carbon fixation can only proceed in this case with electrons deposited into the $NAD^+$/NADH pool. Sulfate reduction begins by the activation of sulfate with ATP forming adenylyl sulfate (APS):[23–25]

$$SO_4^{2-} + ATP + 2H^+ \rightarrow APS + PP_i \qquad (27)$$

Pyrophosphate ($PP_i$) is hydrolyzed to regenerated phosphate according to

$$PP_i + H_2O \rightarrow 2P_i \qquad (28)$$

APS reduction to sulfite is coupled to quinol oxidation, resulting in

$$APS + QH_2 \rightarrow HSO_3^- + AMP + H_{out}^+ \qquad (29)$$

Finally, sulfite is converted to sulfide via a two-step process involving the formation of a trisulfide bond with the protein DsrC with the supply of 4 electrons from the quinone pool and additional two electrons from a cytoplasmic source we assume is NADH.[24] The overall reaction for sulfate reduction is then given by

$$\begin{aligned} &SO_4^{2-} + ATP + 3QH_2 + NADH + 2H_{in}^+ \\ &\quad \rightarrow S^{2-} + AMP + 2P_i + 2Q + NAD^+ + 3H_2O + 3H_{out}^+ \end{aligned} \qquad (30)$$

We assume complex I of the ETC forms reduced quinol from NADH,

$$NADH + 5H_{in}^+ + Q \rightarrow NAD^+ + 4H_{out}^+ + QH_2 \qquad (31)$$

Then, the overall reaction for sulfate reduction is written as

$$\begin{aligned} &4NADH + SO_4^{2-} + ATP + 17H_{in}^+ \\ &\quad \rightarrow 4NAD^+ + S^{2-} + AMP + 2P_i + 3H_2O + 15H_{out}^+ \end{aligned} \qquad (32)$$

Assuming AMP reacts with a second ATP to generate two ADP, both of which are regenerated to ATP, we rewrite the overall reaction for sulfate reduction as

$$4NADH + SO_4^{2-} + 11H_{in}^+ \rightarrow 4NAD^+ + S^{2-} + 3H_2O + 9H_{out}^+ \qquad (33)$$

**Overall stoichiometry**

The preceding analysis allows us to derive the overall stoichiometry of electron-driven $CO_2$-fixation to sucrose for each carbon-fixation pathway (Calvin, rTCA, rGlyP, CETCH, WLP), electron sink (quinone pool, NADH pool), and terminal electron acceptor ($O_2$, $NO_3^-$, $ClO_4^-$, $SO_4^{2-}$), summarized below.

*Calvin cycle*

---

$QH_2$ pool, $O_2$

$$12CO_2 + 121e^- + 18.25O_2 + 121H^+ \rightarrow C_{12}H_{22}O_{11} + 49.5H_2O \quad (34)$$

NADH pool, $O_2$

$$12CO_2 + 75.33e^- + 6.83O_2 + 75.33H^+ \rightarrow C_{12}H_{22}O_{11} + 26.66H_2O \quad (35)$$

$QH_2$ pool, $NO_3^-$

$$12CO_2 + 321.75e^- + 54.75NO_3^- + 376.5H^+ \rightarrow C_{12}H_{22}O_{11} + 23.375N_2 + 177.25H_2O \quad (36)$$

NADH pool, $NO_3^-$

$$12CO_2 + 91.93e^- + 8.79NO_3^- + 100.72H^+ \rightarrow C_{12}H_{22}O_{11} + 4.395N_2 + 39.37H_2O \quad (37)$$

$QH_2$ pool, $ClO_4^-$

$$12CO_2 + 194e^- + 18.25ClO_4^- + 194H^+ \rightarrow C_{12}H_{22}O_{11} + 18.25Cl^- + 86H_2O \quad (38)$$

NADH pool, $ClO_4^-$

$$12CO_2 + 83.14e^- + 4.39ClO_4^- + 83.14H^+ \rightarrow C_{12}H_{22}O_{11} + 4.39Cl^- + 30.56H_2O \quad (39)$$

NADH pool, $SO_4^{2-}$

$$12CO_2 + 157.33e^- + 13.66SO_4^{2-} + 157.33H^+ \rightarrow C_{12}H_{22}O_{11} + 13.66S^{2-} + 67.64H_2O \quad (40)$$

---

*rTCA cycle*

---

$QH_2$ pool, $NO_3^-$

$$12CO_2 + 266.76e^- + 43.75NO_3^- + 310.5H^+ \rightarrow C_{12}H_{22}O_{11} + 21.875N_2 + 144.25H_2O \quad (41)$$

NADH pool, $NO_3^-$

$$12CO_2 + 76.21e^- + 5.64NO_3^- + 81.84H^+ \rightarrow C_{12}H_{22}O_{11} + 2.82N_2 + 29.92H_2O \quad (42)$$

$QH_2$ pool, $ClO_4^-$

$$12CO_2 + 164.66e^- + 14.58ClO_4^- + 164.66H^+ \rightarrow C_{12}H_{22}O_{11} + 14.58Cl^- + 71.32H_2O \quad (43)$$

NADH pool, $ClO_4^-$

$$12CO_2 + 70.57e^- + 2.82ClO_4^- + 70.57H^+ \rightarrow C_{12}H_{22}O_{11} + 2.82Cl^- + 24.28H_2O \quad (44)$$

<u>NADH pool, $SO_4^{2-}$</u>

$$12CO_2 + 118.22e^- + 8.78SO_4^{2-} + 118.22H^+ \rightarrow C_{12}H_{22}O_{11} + 8.78S^{2-} + 48.12H_2O \quad (45)$$

---

*rGlyP*

---

<u>$QH_2$ pool, $O_2$</u>

$$12CO_2 + 101e^- + 13.25O_2 + 101H^+ \rightarrow C_{12}H_{22}O_{11} + 39.5H_2O \quad (46)$$

<u>NADH pool, $O_2$</u>

$$12CO_2 + 62e^- + 3.5O_2 + 62H^+ \rightarrow C_{12}H_{22}O_{11} + 20H_2O \quad (47)$$

<u>$QH_2$ pool, $NO_3^-$</u>

$$12CO_2 + 246.75e^- + 39.75NO_3^- + 286.5H^+ \rightarrow C_{12}H_{22}O_{11} + 19.875N_2 + 132.25H_2O \quad (48)$$

<u>NADH pool, $NO_3^-$</u>

$$12CO_2 + 70.5e^- + 4.5NO_3^- + 75H^+ \rightarrow C_{12}H_{22}O_{11} + 2.25N_2 + 26.5H_2O \quad (49)$$

<u>$QH_2$ pool, $ClO_4^-$</u>

$$12CO_2 + 154e^- + 13.25ClO_4^- + 154H^+ \rightarrow C_{12}H_{22}O_{11} + 13.25Cl^- + 66H_2O \quad (50)$$

<u>NADH pool, $ClO_4^-$</u>

$$12CO_2 + 66e^- + 2.25ClO_4^- + 66H^+ \rightarrow C_{12}H_{22}O_{11} + 2.25Cl^- + 22H_2O \quad (51)$$

<u>NADH pool, $SO_4^{2-}$</u>

$$12CO_2 + 104e^- + 7SO_4^{2-} + 104H^+ \rightarrow C_{12}H_{22}O_{11} + 7S^{2-} + 41H_2O \quad (52)$$

---

*CETCH cycle*

---

<u>$QH_2$ pool, $O_2$</u>

$$12CO_2 + 119.66e^- + 17.92O_2 + 119.66H^+ \rightarrow C_{12}H_{22}O_{11} + 48.84H_2O \quad (53)$$

<u>NADH pool, $O_2$</u>

$$12CO_2 + 72.66e^- + 6.17O_2 + 72.66H^+ \rightarrow C_{12}H_{22}O_{11} + 25.33H_2O \quad (54)$$

<u>$QH_2$ pool, $NO_3^-$</u>

$$12CO_2 + 316.75e^- + 53.75NO_3^- + 370.5H^+ \rightarrow C_{12}H_{22}O_{11} + 26.875N_2 + 174.25H_2O \quad (55)$$

<u>NADH pool, $NO_3^-$</u>

$$12CO_2 + 90.5e^- + 8.5NO_3^- + 99H^+ \rightarrow C_{12}H_{22}O_{11} + 4.25N_2 + 38.5H_2O \quad (56)$$

QH$_2$ pool, $ClO_4^-$

$$12CO_2 + 191.33e^- + 17.92ClO_4^- + 191.33H^+ \rightarrow C_{12}H_{22}O_{11} + 17.92Cl^- + 84.67H_2O \quad (57)$$

NADH pool, $ClO_4^-$

$$12CO_2 + 82e^- + 4.25ClO_4^- + 82H^+ \rightarrow C_{12}H_{22}O_{11} + 4.25Cl^- + 30H_2O \quad (58)$$

NADH pool, $SO_4^{2-}$

$$12CO_2 + 141.33e^- + 9.67SO_4^{2-} + 141.33H^+ + 8FAD \rightarrow C_{12}H_{22}O_{11} + 9.67S^{2-} + 51.68H_2O + 8FADH_2 \quad (59)$$

---

*WLP*

---

QH$_2$ pool, $NO_3^-$

$$12CO_2 + 261.75e^- + 42.75NO_3^- + 304.5H^+ \rightarrow C_{12}H_{22}O_{11} + 21.375N_2 + 141.25H_2O \quad (60)$$

NADH pool, $NO_3^-$

$$12CO_2 + 74.78e^- + 5.36NO_3^- + 80.14H^+ \rightarrow C_{12}H_{22}O_{11} + 2.68N_2 + 29.07H_2O \quad (61)$$

QH$_2$ pool, $ClO_4^-$

$$12CO_2 + 162e^- + 14.25ClO_4^- + 162H^+ \rightarrow C_{12}H_{22}O_{11} + 14.25Cl^- + 70H_2O \quad (62)$$

NADH pool, $ClO_4^-$

$$12CO_2 + 69.43e^- + 2.68ClO_4^- + 69.43H^+ \rightarrow C_{12}H_{22}O_{11} + 2.68Cl^- + 23.72H_2O \quad (63)$$

NADH pool, $SO_4^{2-}$

$$12CO_2 + 114.66e^- + 8.33SO_4^{2-} + 114.66H^+ \rightarrow C_{12}H_{22}O_{11} + 8.33S^{2-} + 46.33H_2O \quad (64)$$

---

**Energy demand and efficiency calculations**

Prior to a more detailed modeling assessment (see Theory section below), preliminary energy demand and efficiency calculations (Table 1) can be determined directly from the stoichiometry. The energy demand $e$ is given by

$$e_k = \frac{n_{e,k} F \Delta V_{0,k}}{M_S} \quad (65)$$

where $n_{e,k}$ is the electron demand per sucrose, $F$ is Faraday's constant, $\Delta V_{0,k}$ is the voltage difference between the anode reaction (water oxidation) and cathode reaction (electron deposition into the quinone pool or NADH pool), and $M_S$ is the molar mass of sucrose for a given physiological mechanism for electron uptake and carbon fixation ($k$). When electrons are deposited into the quinone pool, $\Delta V_0 = 0.918$ V corresponding to the difference between the redox potential of water oxidation (818 mV *vs.* SHE) and electron deposition into electron conduit proteins that interface with the quinone pool (~-100 mV *vs.* SHE). Similarly, $\Delta V_0 = 1.168$ V when electrons are deposited into the NADH pool following the assumption that the electron conduit must accept electrons at a voltage of ~-350 mV *vs.* SHE to reduce $NAD^+$ directly (see eq. 8). The full electrochemical model (see below) accounts for overpotentials associated with electron and chemical species transport and electrochemical kinetics; the energy demand is modified to include these by exchanging $\Delta V_0$ with the $\Delta V$ calculated by the model.

Efficiency is calculated straightforwardly from the combustion energy of sucrose and the energy demand following the method of Claassens *et al.*:[26]

$$\eta_k = \frac{\Delta H^\circ}{M_\mathrm{S} e_k} \tag{66}$$

**Sucrose production from $H_2$**

We rely on a previously described framework for analyzing the production of sucrose from aerobic autotrophic growth on $H_2$.[27] $H_2$ is first oxidized to $H^+$ by soluble and membrane-bound hydrogenases, resulting in

$$\mathrm{H_2 + NAD^+ \rightarrow NADH + H^+} \tag{67}$$

We use the Calvin cycle for carbon fixation and $O_2$ as the terminal electron acceptor. Following our previous analysis of electromicrobial production systems,[27] we assume the $\mathrm{P/O}$ ratio is 2.5 here. Hence, the overall reaction for sucrose production from $H_2$ is given by

$$\mathrm{12CO_2 + 40.4H_2 + 8.2O_2 \rightarrow C_{12}H_{22}O_{11} + 29.4H_2O} \tag{68}$$

**Life Cycle Impact Assessment**

ISO standards 14040 and 14044 were used as guides for the life cycle impact assessments.[28,29] Unless otherwise stated, background life cycle inventories were obtained from the Product Environmental Footprints dataset.[30] Global warming potentials are calculated using the IPCC 2013 100-year model.[31] The functional unit is 1 kg of sucrose at a titer of 100 g/L. The analysis is cradle-to-gate, ending with an unpurified 100 g/L solution of sucrose. However, biologically-sequestered carbon does not count as "negative" emissions at the 100-year time scale. As an example, the overall global warming potential ($C_k$) of the bio-GDE system is given by

$$C_k = e_k c_\mathrm{P} + n_{\mathrm{CO_2}} \frac{M_{\mathrm{CO_2}}}{M_\mathrm{S}} c_\mathrm{DAC} + n_{\mathrm{TEA},k} \frac{M_{\mathrm{TEA},k}}{M_\mathrm{S}} c_\mathrm{TEA} + a_{\mathrm{E},k} \ddot{c}_\mathrm{E} \tag{69}$$

where $c_\mathrm{P}$, $c_\mathrm{DAC}$, and $c_\mathrm{TEA}$ are the carbon footprints of power generation, direct air capture of $CO_2$, and terminal electron acceptor, respectively; $n_{\mathrm{CO_2}}$ and $n_\mathrm{TEA}$ are the demands of $CO_2$ and the terminal electron acceptor per sucrose molecule from stoichiometry; $a_{\mathrm{E},k}$ is the effective area of electrolyzer necessary to produce a unit of product, given by

$$a_{\mathrm{E},k} = \frac{1}{M_\mathrm{S} \gamma t} \left( \frac{n_{\mathrm{e},k} F}{i} \right) \tag{70}$$

and $\ddot{c}_\mathrm{E}$ is the carbon footprint of electrolyzer materials normalized to the electrolyzer area:

$$\ddot{c}_{\mathrm{E}} = \sum_{n} c_n \ddot{m}_n \tag{71}$$

Here, $\gamma$ is the operation uptime, $t$ is the electrolyzer lifetime, $i$ is the bio-GDE current density, $c_n$ is the carbon footprint of electrolyzer material $n$, and $\ddot{m}_n$ is the areal mass (mass per unit area of electrolyzer) of electrolyzer material $n$. Carbon footprints are calculated as described below.

*Terminal Electron Acceptor (Re-generation)*

Upstream production was modeled for four terminal electron acceptors: oxygen, nitrate, sulfate, and perchlorate. Mass and energy balances, along with data from industrial and academic literature, were used to determine material and energy flows for upstream production of each terminal electron acceptor. Oxygen is assumed to be abundant and readily available, so no additional energy or greenhouse gas emissions were included to produce oxygen. In our process model, nitrate is produced from atmospheric nitrogen gas through a combination of the Haber Bosch process and the Ostwald Process.[32] Hydrogen gas for the Haber Bosch process is produced through electrolysis of water driven by wind power; both energy and life cycle greenhouse gas emissions for production of ammonia through this process are obtained from Singh et al.[33] Nitric acid is produced from the combustion of this ammonia through the Ostwald process. As this process is exothermic, no additional energy is required, though nitrous oxide as a byproduct is produced (8 kg $N_2O$/ton $HNO_3$).[34] Our model assumes 95% captured and the balance emitted to the atmosphere.[34] Nitric acid is neutralized by the addition of sodium hydroxide, producing neutral sodium nitrate for the MES process. Perchlorate is produced using the chloride ions generated through perchlorate respiration, forming a chlorine cycle for the process. Sodium chloride is first converted to sodium chlorate by electrolysis.[35] Sodium chlorate is further converted to sodium perchlorate, again by electrolysis.[35] Sulfur is likewise recycled in the process. Hydrogen sulfide gas produced by the sulfate-reducing bacteria is converted to sulfuric acid through the wet sulfuric acid process, an exothermic process which requires no additional energy.[36] Sulfuric acid is neutralized to sodium sulfate by the addition of sodium hydroxide. Sodium hydroxide (in both sodium nitrate and sodium sulfate production) is produced from the electrolysis of sodium chloride and water in the chlor-alkali process, with mass allocation used to determine the impacts associated with NaOH production. Electricity demand is assumed to be the major contributor of the global warming potential of the chlor-alkali process.[37]

*Other LCA Components*

All electricity in the process is assumed to be from wind energy, with life cycle impacts drawn from the PEF dataset. It should be noted other decarbonized electricity sources such as thin-film photovoltaics and hydropower have similar global warming potentials and would therefore lead to similar results. Carbon dioxide in the modelled process is provided via direct air capture, with energy requirements and life cycle impacts obtained from Duetz and Bardow.[38] Impacts for most of electrolyzer materials ($IrO_2$, $SnO_2$, carbon paper, and PMMA) are obtained from the PEF database while impacts for Nafion are obtained from Stropnik et al.[39] Electrolyzer material requirements are calculated using the system described by Xu *et al.*[40] as a basis assuming a geometric area of 1 $m^2$, an electrode separation distance of 1 cm, and an electrolyzer lifetime of three years assumed. In the case of direct electroautotrophy, the bio-GDE uses the geometry and density of a Sigracet 35BC gas diffusion electrode for the cathode and makes the same assumptions regarding geometric area, electrode separation, and lifetime.

Electroautotrophic production of sucrose is compared to two alternative sucrose production methods: from sugarcane and in a metabolically engineered Knallgas bacterium. Life cycle global warming potential of sugarcane is derived from Izursa *et al.*,[41] while that of sucrose production in Knallgas bacterium is determined by adapting the process model from our previous work.[27]

### Theory

#### Mass transport in a bio-GDE

*Governing equations*

Following Weng *et al.*[42], we describe the gas diffusion electrode with a microbial catalyst layer (bio-GDE) using a one-dimensional, macro-homogeneous model assuming isothermal conditions. We explicitly consider both gas and liquid phases, species transport within each phase, and mass transfer between the two phases. With this model, we were attempting only to determine if gas-liquid mass transfer or gas-phase transport could limit the productivity of a bio-GDE. Therefore, we considered only two species: $CO_2$ and $O_2$, although $N_2$ was included in the gas phase at a small concentration (1000 ppm) for numerical stability. The mole balance within the liquid phase of the bio-GDE can be written as

$$\frac{\partial c_i^{\mathrm{L}}}{\partial t} = -\frac{\partial n_i^{\mathrm{L}}}{\partial x} - R_{\mathrm{X},i} + R_{\mathrm{GL},i} \tag{72}$$

where $c_i^{\mathrm{L}}$ is the liquid phase concentration, $n_i^{\mathrm{L}}$ is the molar flux in the liquid phase, $R_{\mathrm{X},i}$ and $R_{\mathrm{GL},i}$ are the microbial reaction (carbon fixation) and gas-liquid mass transfer source terms, respectively. The subscript $i$ refers to the chemical species. In the gas phase, the mole balance is given by

$$\frac{\partial c_i^{\mathrm{G}}}{\partial t} = -\frac{\partial n_i^{\mathrm{G}}}{\partial x} - \frac{\theta^{\mathrm{L}}}{\theta^{\mathrm{G}}} R_{\mathrm{GL},i} \tag{73}$$

Where $\theta^{\mathrm{L}}$ and $\theta^{\mathrm{G}}$ are the liquid- and gas-phase volume fractions, respectively. Note that the gas-liquid mass transfer term, $R_{\mathrm{GL},i}$ is written as positive for the liquid phase (*i.e.,* a source term) and negative for the gas-phase (*i.e.,* a sink term).

*Geometry*

We assume that the biocatalyst layer (bCL) within the bio-GDE has a defined volume fraction for the conductive support material ($\theta^{\mathrm{CM}}$) with a defined specific surface area ($a_{\mathrm{v}}^{\mathrm{CM}}$). If the biomass is distributed evenly throughout the bCL, the volume fraction of biomass is given by

$$\theta^{\mathrm{X}} = \gamma t^{\mathrm{X}} a_{\mathrm{v}}^{\mathrm{CM}} \tag{74}$$

where $\theta^{\mathrm{X}}$ is the biomass volume fraction, $\gamma$ is the packing factor, and $t^{\mathrm{X}}$ is the biofilm thickness. We assume $\gamma \approx 0.52$, equivalent to a square lattice of spheres with equal diameter. The surface area of the biomass is written as

$$a_{\mathrm{v}}^{\mathrm{X}} = 6\gamma a_{\mathrm{v}}^{\mathrm{CM}} \tag{75}$$

To determine the volume fraction of liquid and gas phases in the bCL, we define a saturation fraction ($S$) such that

$$\theta^{\mathrm{L}} = S\left(1 - \theta^{\mathrm{CM}} - \theta^{\mathrm{X}}\right) \tag{76}$$

and

$$\theta^{\mathrm{G}} = 1 - \theta^{\mathrm{CM}} - \theta^{\mathrm{X}} - \theta^{\mathrm{L}} \tag{77}$$

We use $S = 0.64$ following the "ideally wetted" case in Weng *et al.*[42] To calculate the liquid film thickness (which is relevant for gas-liquid mass transfer as discussed later), we use

$$t^{\mathrm{L}} = \frac{\theta^{\mathrm{L}}}{a_{\mathrm{v}}^{\mathrm{X}}} \tag{78}$$

Finally, the specific surface area of the liquid phase is given by

$$a_{\mathrm{v}}^{\mathrm{L}} = \frac{6\theta^{\mathrm{X}}}{t^{\mathrm{X}} + t^{\mathrm{L}}} \tag{79}$$

following geometric arguments.

*Liquid-phase transport*

Flux of aqueous species is given simply by

$$n_i^{\mathrm{L}} = -D_i^{\mathrm{L,eff}} \frac{\partial c_i^{\mathrm{L}}}{\partial x} \tag{80}$$

where $D_i^{\mathrm{L,eff}}$ is the effective diffusion coefficient of species $i$ in the liquid phase. Because species transport is occurring through the porous bCL, we use effective diffusion coefficients following the Bruggeman relationship,

$$D_i^{\mathrm{L,eff}} = \frac{\theta^{\mathrm{L}}}{\tau^{\mathrm{L}}} D_i^{\mathrm{L}} = \left(\theta^{\mathrm{L}}\right)^{3/2} D_i^{\mathrm{L}} \tag{81}$$

where $\tau^{\mathrm{L}}$ is the tortuosity of the medium. Our full electrochemical model, as described in the next section, uses the Nernst-Planck relationship to describe both diffusion and migration in the liquid phase. We neglect migration here because none of the species we consider ($CO_2$, $O_2$, $N_2$) are charged.

*Gas-phase transport*

Gaseous species flux consists of both diffusive and convective terms,

$$n_i^{\mathrm{G}} = M_i j_i^{\mathrm{G}} + M_i \rho_i u_{\mathrm{g}} \tag{82}$$

where $M_i$ is the molar mass, $j_i$ is the diffusive mass flux, $\rho_i$ is the mass density, and $u_{\mathrm{g}}$ is the mass-averaged fluid velocity. Following Weng *et al.*,[42] the diffusive flux is calculated using a mixture-averaged diffusion model,

$$j_i^{\mathrm{G}} = -\rho_{\mathrm{g}} D_i^{\mathrm{G,eff}} \frac{\partial \omega_i}{\partial x} - \rho_{\mathrm{g}} D_i^{\mathrm{G,eff}} \omega_i \frac{\partial M_n}{M_n \partial x} \tag{83}$$

where $\omega_i$ is the mass fraction of species $i$, $\rho_\mathrm{g}$ is the gaseous mixture density, $M_\mathrm{n}$ is the average molar mass of the mixture, $M_\mathrm{n} = \left(\sum_i \frac{\omega_i}{M_i}\right)^{-1}$, and $D_i^\mathrm{G,eff}$ is the effective gas-phase diffusion coefficient for species $i$. The gas-phase diffusion coefficient includes a mass-averaged Stefan-Maxwell diffusivity ($D_i^\mathrm{m}$) and Knudsen diffusivity ($D_i^\mathrm{K}$) occurring in parallel:

$$D_i^\mathrm{G} = \left(\frac{1}{D_i^\mathrm{m}} + \frac{1}{D_I^\mathrm{K}}\right)^{-1} \tag{84}$$

with

$$D_i^\mathrm{m} = \frac{1-\omega_i}{\sum_{n\neq i}\frac{y_n}{D_{i,n}}} \tag{85}$$

and

$$D_i^\mathrm{K} = \frac{2r_\mathrm{p}}{3}\sqrt{\frac{8RT}{\pi M_i}} \tag{86}$$

Here, $r_\mathrm{p}$ is the pore radius of the porous medium, $y_i$ is the molar fraction of species $i$, $R$ is the gas constant and $T$ is the temperature. We also correct the diffusivity in the gas phase to account for porosity and tortuosity:

$$D_i^\mathrm{G,eff} = \frac{\theta^\mathrm{G}}{\tau^\mathrm{G}} D_i^\mathrm{G} = \left(\theta^\mathrm{G}\right)^{3/2} D_i^\mathrm{G} \tag{87}$$

The mass-averaged velocity field ($u_\mathrm{g}$) is described using Darcy's law,

$$u_g = -\frac{\kappa^\mathrm{G}}{\mu^\mathrm{G}}\frac{\partial p}{\partial x} \tag{88}$$

Where $\kappa^\mathrm{G}$ is the permeability, $\mu^\mathrm{G}$ is the gas-phase viscosity, and $p$ is the total gas pressure. The $N$th gas species mass fraction is determined by a mass balance:

$$\sum_i \omega_i = 1 \tag{89}$$

*Carbon fixation reaction*

Carbon fixation relies on the enzymatic conversion of $CO_2$ into metabolites (*e.g.*, pyruvate, acetyl-CoA, *etc.*) and requires energy in the form of ATP, NADH, and/or $Fd^{2-}$, which are produced by electron uptake and (in the case we consider in this model) aerobic respiration. Hence, either the carbon fixation pathway or aerobic respiration can set a limit on the overall rate of carbon fixation. We calculated the enzymatic rate limits using Michaelis-Menten kinetics:

$$r_{\mathrm{CO_2}} = k_{\mathrm{cat,CO_2}} E_{\mathrm{CO_2}} \left(\frac{c_{\mathrm{CO_2}}^\mathrm{L}}{K_{\mathrm{M,CO_2}} + c_{\mathrm{CO_2}}^\mathrm{L}}\right) \tag{90}$$

$$r_{O_2} = \alpha_n k_{\mathrm{cat,O_2}} E_{O_2} \left( \frac{c_{O_2}^{\mathrm{L}}}{K_{\mathrm{M,O_2}} + c_{O_2}^{\mathrm{L}}} \right)$$

where $\alpha_n$ is a stoichiometric coefficient depending on the ratio of $O_2$ to $CO_2$ consumed during carbon-fixing metabolism, $k_{\mathrm{cat},i}$ is the turnover number of the relevant enzyme, $E_i$ is the intracellular enzyme concentration, and $K_{\mathrm{M},i}$ is the Michaelis constant. We set $k_{\mathrm{cat,CO_2}}$ equal to 100 $s^{-1}$ for carbon fixation using the reductive glycine pathway (rGlyP) to represent an upper bound in accordance with the maximum rate of formate dehydrogenases in *Cupriavidus necator* that would be responsible for the first carbon fixation step in the rGlyP,[43] and used an intracellular enzyme concentration of ~220 mM based on the estimate that RuBisCo comprises 3% of the enzymes in carbon-fixing organisms using the Calvin cycle.[44,45] For $O_2$ respiration, we calculated the product $k_{\mathrm{cat,O_2}} E_{O_2}$ based on estimates of cellular respiration rates in *E. coli*.[46] In all cases we considered in our bio-GDE model, the carbon fixation pathway set a more stringent bound on the rate of carbon fixation than the aerobic respiration pathway (*i.e.*, $r_{CO_2} < r_{O_2}$ under all conditions considered in our model). Hence, the microbial carbon fixation reaction is written as

$$R_{\mathrm{X},i} = \alpha_i \theta^{\mathrm{X}} r_{CO_2} \tag{91}$$

where $\alpha_i$ is a stoichiometric coefficient ($\alpha_{CO_2}$ is always equal to 1, but $\alpha_{O_2}$ varies depending on the electron sink and carbon fixation pathway, as discussed in the section on the metabolism of carbon fixation).

*Gas-liquid mass transfer*

Gas-phase $CO_2$ and $O_2$ dissolves into the liquid at the gas/liquid interface. Following Weng *et al.*,[42] the gas-liquid mass transfer coefficient ($k_{\mathrm{GL},i}$) is calculated according to

$$k_{\mathrm{GL},i} = \frac{D_i^{\mathrm{L}}}{t^{\mathrm{L}}} \tag{92}$$

Then, the rate of gas-liquid mass transfer is given by

$$R_{\mathrm{GL},i} = a_{\mathrm{v}}^{\mathrm{L}} k_{\mathrm{GL},i} \left( H_i p_{\mathrm{G}} y_i - c_i^{\mathrm{L}} \right) \tag{93}$$

where $H_i$ is Henry's constant for species $i$.

*Boundary conditions*

In the liquid phase, we use no-flux boundary conditions at the electrolyte-bCL and bCL-gas diffusion layer (GDL) interfaces. In the latter case, this is because the liquid phase ends at the bCL-GDL interface, so no liquid-phase flux is possible. In the former case, this is because the rate of diffusion of sparingly soluble gases from the bulk electrolyte, through the fluid boundary layer, to the electrolyte-bCL boundary is several orders of magnitude lower than the rate of gas-liquid mass transfer within the bCL such that this diffusive contribution to the liquid-phase concentration of $CO_2$ and $O_2$ can be neglected.

For the gas-phase, we use a no-flux boundary condition at the electrolyte-bCL interface because, although gas-liquid mass transfer would occur, the surface area of the bCL-electrolyte interface is several orders of magnitude smaller than the gas-liquid interfacial area within the bCL. At the bCL-GDL interface, the gas feed composition is set to 50 mol% $CO_2$, 49.9 mol% $O_2$, and 0.1 mol% $N_2$.

*Model analysis*

We calculated the projected surface area current density as

$$i = \int_0^{l_{\mathrm{bCL}}} n_{\mathrm{e}} F R_{\mathrm{X,CO_2}} \partial x \tag{94}$$

where $l_{\mathrm{bCL}}$ is the biocatalyst layer thickness, $n_{\mathrm{e}}$ is the stoichiometric ratio of electrons consumed per $CO_2$ fixed, and F is Faraday's constant.

**Full electrochemical bio-GDE model**

*System overview*

The model considers a one-dimension bio-electrochemical reactor for the conversion of $CO_2$ into sucrose via direct electron transfer in a bio-GDE. The reactor has a well-mixed region that is exchanged at a fixed dilution rate and to which a $CO_2$-containing gas mixture is constantly supplied at a fixed pressure. Fluid boundary layers (BLs) separate the well-mixed liquid phase from the anode surface and the bio-GDE. Electrochemical reactions at the anode surface (water oxidation) and throughout the biocatalyst layer (bCL) within the bio-GDE (direct electron transfer coupled to $CO_2$-fixation) are driven by an applied voltage. Microbes attached to the conductive support material (CM) are assumed to accept electrons into the quinone or NADH pool, use $O_2$ as the terminal electron acceptor, and use the reductive glycine pathway (rGlyP) to fix $CO_2$. The chemical species we consider in the reactor system are dissolved $CO_2$, bicarbonate anions ($HCO_3^-$), carbonate anions ($CO_3^{2-}$), protons ($H^+$), hydroxide anions ($OH^-$), sodium cations ($Na^+$), and nitrate anions ($NO_3^-$). $NO_3^-$ was selected as a representative anion for sodium salt to avoid the use of chloride ions ($Cl^-$), which are known to produce deleterious and toxic side reactions at the cathode surface in MES systems.[47] We don't consider dissolved $O_2$ in the model because the previous bio-GDE model demonstrated that the bio-GDE enables saturated $O_2$ throughout the liquid phase even at current densities >170 mA/cm$^2$ with NADH as the electron sink and >280 mA/cm$^2$ with $QH_2$ as the electron sink.

We assume that the microbes within the bCL are not actively growing and are instead acting as a compartment for enzymatic processes. Hence, we assume that all fixed carbon is diverted to sucrose, and that additional nutrients (*e.g.*, ammonia, phosphates, *etc.*) are unnecessary. We emphasize that these assumptions do not correspond to any experimentally-realized system; rather, our goal here is to determine the best-case productivity and efficiency of a DET-based electromicrobial production system and the operating conditions of such a system. We also neglect cellular maintenance requirements, which have been previously shown to be negligible for calculations of this type.[48]

*Well-mixed phase balance equations*

The well-mixed electrolyte region is assumed to have sufficient convective mixing such that no concentration gradients are formed. The well-mixed phase mole balance is given for our reactor model by

$$\frac{dc_i}{dt} = R_{\mathrm{A-B},i} + R_{\mathrm{F},i} + R_{\mathrm{GL},i} + S_{\mathrm{A}}\left(n_i|_{\mathrm{BL_A}} - n_i|_{\mathrm{BL_C}}\right) \tag{95}$$

where $c_i$ is the concentration, $R_i$ is the net volumetric rate of formation and consumption due to acid-base reactions (A-B), electrolyte flow (F), and gas-liquid mass transfer (GL), and $n_i$ is the flux of species $i$. The projected electrode surface area-to-volume ratio is given by $S_{\mathrm{A}}$. By convention, the positive xdirection is defined from left to right such that species flux from the cathode boundary layer phase ($\mathrm{BL_C}$) to the well-mixed phase will have a negative value.

*Anode and cathode boundary layer balance equations*

Fluid in the boundary layers is approximately stagnant. Hence, species transport in these regions occurs via by diffusion and migration driven by electrochemical potential gradients. The mole balances in both the anode and cathode BL phases are given by

$$\frac{\partial c_i}{\partial t} = -\frac{\partial n_i}{\partial x} + R_{\mathrm{A-B},i} \tag{96}$$

since neither electrolyte flow nor gas-liquid mass transfer occurs here.

*Biocatalyst layer balance equation*

In the biocatalyst layer, the $CO_2$-fixation reaction and gas-liquid mass transfer provide additional sink/source terms for species in the liquid phase. Hence, the balance equation is written as

$$\frac{\partial c_i}{\partial t} = -\frac{\partial n_i}{\partial x} + R_{\mathrm{A-B},i} + R_{\mathrm{GL},i}^{\mathrm{bCL}} + R_{\mathrm{X},i} \tag{97}$$

where $R_{\mathrm{GL},i}^{\mathrm{bCL}}$ is the gas-liquid mass transfer term in the bCL phase and $R_{\mathrm{X},i}$ accounts for the $CO_2$-fixation reaction.

*Species transport in the boundary layers and biocatalyst layer*

The molar flux of species in dilute electrolyte solutions is written as the sum of diffusive and migrative fluxes:

$$n_i = -D_i^{\mathrm{eff}} \frac{\partial c_i}{\partial x} - z_i u_i F c_i \frac{\partial \phi_1}{\partial x} \tag{98}$$

where $D_i^{\mathrm{eff}}$ and $u_i$ are the diffusivity and mobility (related by the Nernst-Einstein relationship, $u_i = D_i/RT$ for dilute solutions) of species $i$, $z_i$ is the charge number, $F$ is Faraday's constant, and $\phi_1$ is the local electrolyte potential. Effective diffusion coefficients are used in the bCL according to the Bruggeman relationship as described for the bio-GDE in the preceding section. The net ionic current density in the electrolyte ($i_1$) can be calculated from the total ionic flux:

$$i_1 = F \sum_i z_i n_i \tag{99}$$

following electroneutrality,

$$\sum_i z_i c_i = 0 \tag{100}$$

*Acid-base reactions*

The acid-base carbon dioxide/bicarbonate/carbonate and water dissociation reactions occur in each liquid phase and are treated as kinetic expressions without assuming equilibrium:

$$CO_{2(aq)} + H_2O \overset{k_{+1},k_{-1}}{\longleftrightarrow} H^+ + HCO_3^- \qquad K_1 \qquad (101)$$

$$HCO_3^- \overset{k_{+2},k_{-2}}{\longleftrightarrow} H^+ + CO_3^{2-} \qquad K_2 \qquad (102)$$

$$CO_{2(aq)} + OH^- \overset{k_{+3},k_{-3}}{\longleftrightarrow} HCO_3^- \qquad K_3 = K_1/K_W \qquad (103)$$

$$HCO_3^- + OH^- \overset{k_{+4},k_{-4}}{\longleftrightarrow} CO_3^{2-} + H_2O \qquad K_4 = K_2/K_W \qquad (104)$$

$$H_2O \overset{k_W,k_{-W}}{\longleftrightarrow} H^+ + OH^- \qquad K_W \qquad (105)$$

where $k_{+n}$ and $k_{-n}$ are the forward and reverse rate constants, respectively, and $K_n$ is the equilibrium constant for the $n$th reaction, given by

$$K_n = \exp\left(\frac{\Delta S_n}{R}\right)\exp\left(-\frac{\Delta H_n}{RT}\right) \qquad (106)$$

where $\Delta S_n$ and $\Delta H_n$ are the molar entropy change and heat of reaction, respectively, for reaction $n$. Source and sink terms resulting from these reactions are compiled in $R_{A-B,i}$, written as

$$R_{A-B,i} = \sum_n \nu_i \left( k_{+n} \prod_{\nu_i<0} c_i - k_{-n} \prod_{\nu_i>0} c_i \right) \qquad (107)$$

where $\nu_i$ is the stoichiometric coefficient of species $i$ for the $n$th reaction and reverse rate constants are calculated from:

$$k_{-n} = \frac{k_{+n}}{K_n} \qquad (108)$$

*Electrolyte flow*

Liquid electrolyte is fed to and extracted from the well-mixed liquid phase at a constant dilution rate, resulting in a feed term written as

$$R_{F,i} = D_{liq}(c_{F,i} - c_i) \qquad (109)$$

where $D_{liq}$ is the liquid dilution rate and $c_{F,i}$ is the feed concentration of species $i$. For the life cycle analysis, we defined the functional unit of analysis to be sucrose at 100 g/L. We therefore defined the dilution rate as a function of the projected surface area current density to maintain a sucrose titer of 100 g/L in the well-mixed phase, given by:

$$D_{liq} = \frac{i S_A M_S}{n_e F c'_S} \qquad (110)$$

where $i$ is the projected surface area current density, $M_\mathrm{S}$ is the molar mass of sucrose (342.3 g/mol), $n_\mathrm{e}$ is the number of electrons consumed per sucrose molecule, and $c'_\mathrm{S}$ is the desired sucrose mass concentration (100 g/L in our case).

*Gas feed*

A $CO_2$-containing gas mixture is fed into the well-mixed liquid phase at a total pressure $P$, resulting in mass transfer according to

$$R_{\mathrm{GL,CO_2}} = k_{\mathrm{L,CO_2}} a\left(P\beta_{\mathrm{CO_2}} y_{\mathrm{F,CO_2}} - c_{\mathrm{CO_2}}\right) \tag{111}$$

where $k_{\mathrm{L,CO_2}}a$ is the volumetric mass transfer coefficient on the liquid side of the gas/liquid interface, $\beta_{\mathrm{CO_2}}$ is the Bunsen solubility coefficient, and $y_{\mathrm{F,CO_2}}$ is the mole fraction of $CO_2$ in the gas phase. The Bunsen solubility coefficient, $\beta_{\mathrm{CO_2}}$, is calculated according to

$$\ln\beta = A_1 + A_2\left(\frac{100}{T}\right) + A_3 \ln\left(\frac{T}{100}\right) + S\left[B_1 + B_2\left(\frac{T}{100}\right) + B_3\left(\frac{T}{100}\right)^2\right] \tag{112}$$

where $A_n$ and $B_n$ are fitting parameters and $S$ is the salinity (in g/kg). The gas-liquid mass transfer in the bCL is conceptually similar, but the mass transfer coefficient is adjusted to reflect the gas-liquid interfacial area as described in the preceding section:

$$R_{\mathrm{GL,CO_2}}^{\mathrm{bCL}} = a_\mathrm{v}^\mathrm{L} k_{\mathrm{GL,CO_2}}\left(P\beta_{\mathrm{CO_2}} y_{\mathrm{F,CO_2}} - c_{\mathrm{CO_2}}\right) \tag{113}$$

*Electrode reactions – anode*

The surface reaction at the anode is the oxidation of water:

$$\mathrm{H_2O} \longrightarrow \frac{1}{2}\mathrm{O_2} + 2\mathrm{H^+} + 2\mathrm{e^-} \qquad E^0_{\mathrm{OER}} \tag{114}$$

where $E^0_{\mathrm{OER}}$ is the equilibrium potential of the oxygen evolution half-cell reaction (OER) at standard state. The anode reaction is related to species transport by a flux boundary condition at the electrode surface,

$$n_i|_\mathrm{A} = \frac{\nu_i i_\mathrm{R}}{nF} \tag{115}$$

where $i_\mathrm{R}$ is the reaction current density and $n$ is the number of electrons participating in the electrode reaction. We model charge transfer kinetics at the anode using Butler-Volmer kinetics:

$$i_\mathrm{R} = i_0\left[\left(\frac{c_{\mathrm{red}}}{c_{\mathrm{red,0}}}\right)^{\gamma_{\mathrm{red}}} \exp\left(\frac{\alpha_\mathrm{a} F\eta}{RT}\right) - \left(\frac{c_{\mathrm{ox}}}{c_{\mathrm{ox,0}}}\right)^{\gamma_{\mathrm{ox}}} \exp\left(\frac{\alpha_\mathrm{c} F\eta}{RT}\right)\right] \tag{116}$$

where $i_0$ is the exchange current density, $\gamma_{\mathrm{red/ox}}$ is the reaction order with respect to a reactant, $\alpha_{\mathrm{a/c}}$ is the anodic/cathodic transfer coefficient, and $\eta$ is the overpotential. The overpotential is defined according to

$$\eta = \phi_s - \phi_l - E \tag{117}$$

where $\phi_s$ is the electrode potential, $\phi_l$ is the electrolyte potential, and $E$ is the half-cell equilibrium potential.

Because water oxidation creates acidic conditions near the anode surface, bicarbonate and carbonate species will be converted to aqueous $CO_2$ according to Le Chatelier's principle. To avoid the unrealistic supersaturation this would cause, we describe the evolution of $CO_2$ as an additional sink term for $CO_2$ in the electrolyte as

$$R_{evo,CO_2} = \begin{cases} -\gamma_{CO_2} S_{CO_2}^2 & S \geq 1 \\ 0 & S < 1 \end{cases} \tag{118}$$

where $\gamma_{CO_2}$ is the releasing coefficient and $S_{CO_2}$ is the supersaturation coefficient, defined as $c_{CO_2}/\beta_{CO_2} p_{CO_2}$, where $p_{CO_2}$ is the partial pressure.[49–52]

*Electrode reactions – biocatalyst layer*

The $CO_2$-fixation reaction ($R_{X,i}$) is related to the current density in the bCL by

$$R_{X,i} = \frac{\nu_i a_v^X i_X}{n_e F} \tag{119}$$

where $a_v^X$ is the active specific surface area of the microbes in the bCL and $i_X$ is the current density on the microbial surfaces. The active specific surface area was calculated in the preceding section. The current density, and therefore the $CO_2$-fixation rate, can be limited either by the kinetics of electron transfer or by the enzymatic rate limit of $CO_2$ fixation. To account for this, we describe $i_X$ as

$$i_X = \frac{i_R}{1 + \left| \frac{i_R}{i_{lim}} \right|} \tag{120}$$

where $i_R$ is the reaction current density limit described by Butler-Volmer kinetics,

$$i_R = i_0 \left[ \left( \frac{c_{red}}{c_{red,0}} \right)^{\gamma_{red}} \exp\left( \frac{\alpha_a F \eta}{RT} \right) - \left( \frac{c_{ox}}{c_{ox,0}} \right)^{\gamma_{ox}} \exp\left( \frac{\alpha_c F \eta}{RT} \right) \right] \tag{121}$$

and $i_{lim}$ is the biomass-limited current density. We calculate the biomass-limited current density by projecting the enzymatic $CO_2$-fixation rate limit to the total cell surface:

$$i_{lim} = n_e F k_{cat} E_{CO_2} \frac{\theta^X}{a_v^X} \tag{122}$$

where $k_{cat}$ is the enzyme turnover number and $E_{CO_2}$ is the intracellular concentration of the rate-limiting enzyme in the $CO_2$-fixation pathway. This formulation relies on the fact that intracellular substrate diffusion is much faster than rate-limiting enzymatic reaction steps in carbon fixation pathways.[50]

*Electron transport in solid electrodes*

Electron transport in the solid electrode (the anode and the conductive support material within the bCL) is governed by charge conservation and Ohm's law, given by:

$$\nabla i_{\mathrm{s}} = -\nabla i_{\mathrm{l}} = -a_{\mathrm{v}}^{\mathrm{X}} i_{\mathrm{X}} \tag{123}$$

$$i_{\mathrm{s}} = \kappa_{\mathrm{s}} \frac{\partial \phi_{\mathrm{s}}}{\partial x} \tag{124}$$

where $i_{\mathrm{s}}$ is the electrode current density and $\kappa_{\mathrm{s}}$ is the conductivity. We modify the conductivity in the bCL with a Bruggeman correction factor:

$$\kappa_{\mathrm{s}}^{\mathrm{eff}} = \left(\theta^{\mathrm{CM}}\right)^{1.5} \kappa_{\mathrm{s}} \tag{125}$$

**Numerical method**

The equations for both the bio-GDE model and the electrochemical system model are solved using the MUMPS general solver in COMSOL Multiphysics 5.4. For the bio-GDE, the modeling domain has a maximum element size of 0.02 μm. For the electrochemical system, the modeling domain has a maximum element size of 10 μm in the well-mixed regions and 0.5 μm in the boundary layers and bCL to capture steep concentration gradients. Model parameters for the bio-GDE and electrochemical system are listed in Tables S1 and S2, respectively.

**Table S1. Model parameters for the bio-GDE.**

| Parameter | Value | Units | References |
|---|---|---|---|
| *Operating conditions* | | | |
| $T$ | 310.15 | K | fixed |
| $P_0$ | 2 | atm | fixed |
| $y_{\mathrm{f,CO_2}}$ | 0.5 | -- | fixed |
| $y_{\mathrm{f,O_2}}$ | 0.5 | -- | fixed |
| $l_{\mathrm{bCL}}$ | 325 | μm | fixed |
| *Geometry* | | | |
| $\theta^{\mathrm{CM}}$ | 0.2 | -- | 53 |
| $a_{\mathrm{v}}^{\mathrm{CM}}$ | $1\times10^6$ (high surface area)<br>$5.6\times10^5$ (thick biofilm) | $m^{-1}$ | assumed |
| $t^{\mathrm{X}}$ | 1 (high surface area)<br>10 (thick biofilm) | μm | assumed |
| $S$ | 0.64 | -- | 42 |
| $r_{\mathrm{P}}$ | 1.47 | μm | 42 |
| *Liquid-phase diffusion coefficients* | | | |
| $D_{\mathrm{CO_2}}^{\mathrm{L}}$ | $14.6836 \times 10^{-9} \left(\frac{T}{217.2056} - 1\right)^{1.997}$ | $m^2\ s^{-1}$ | 54 |
| $D_{\mathrm{O_2}}^{\mathrm{L}}$ | $10\hat{}\left(8.410 + \frac{773.8}{T} - \left(\frac{506.5}{T}\right)^2\right)$ | $m^2\ s^{-1}$ | 55 |
| *Gas-phase diffusion coefficients* | | | |
| $D_{\mathrm{O_2-CO_2}}$ | 0.156 | $cm^2\ s^{-1}$ | 56 |
| $D_{\mathrm{N_2-CO_2}}$ | 0.165 | $cm^2\ s^{-1}$ | 56 |
| $D_{\mathrm{O_2-N_2}}$ | 0.225 | $cm^2\ s^{-1}$ | 56 |
| *Gas-phase transport parameters* | | | |

| | | | |
|---|---|---|---|
| $\mu^{G}$ | 1.9×10$^{-5}$ | Pa s | [57] |
| $\kappa^{G}$ | 1.72×10$^{-7}$ | cm$^2$ | [42] |
| *Enzyme kinetics* | | | |
| $k_{cat,CO_2}$ | 100 | s$^{-1}$ | [58] |
| $E_{CO_2}$ | 0.224 | mM | Note 1 |
| $K_{M,CO_2}$ | 3.3 | mM | [49] |
| $k_{cat,O_2} \times E_{O_2}$ | 2083 | mmol L$^{-1}$ s$^{-1}$ | [46] |
| $K_{M,O_2}$ | 3 | nM | [59] |

**Table S2. Full electrochemical model parameters.**

| Parameter | Value | Unit | References |
|---|---|---|---|
| *Operating conditions* | | | |
| $T$ | 310.15 | K | fixed |
| $P$ | 2 | atm | fixed |
| $y_{f,CO_2}$ | 0.5 | -- | fixed |
| $c'_{sucrose}$ | 100 | g L$^{-1}$ | fixed |
| *Geometry* | | | |
| $S_A$ | 100 | m$^{-1}$ | assumed |
| $l_{membrane}$ | 100 | μm | assumed |
| $l_{BL}$ | 100 | μm | assumed |
| $l_{bCL}$ | 325 | μm | assumed |
| *Diffusion coefficients* | | | |
| $D_{H^+}$ | $1.56 \times 10^{-10}(T - 273.15) + 5.49 \times 10^{-9}$ | m$^2$ s$^{-1}$ | [49] |
| $D_{OH^-}$ | $4.52 \times 10^{-4} \exp\left(-1618\left(\frac{1}{T} + \frac{1}{273.15}\right)\right)$ | m$^2$ s$^{-1}$ | [49] |
| $D_{Na^+}$ | $8.85 \times 10^{-12} T$ | m$^2$ s$^{-1}$ | [49] |
| $D_{HCO_3^-}$ | $7.016 \times 10^{-9}\left(\frac{T}{204.028} - 1\right)^{2.3942}$ | m$^2$ s$^{-1}$ | [54] |
| $D_{CO_3^{2-}}$ | $5.447 \times 10^{-9}\left(\frac{T}{210.265} - 1\right)^{2.192}$ | m$^2$ s$^{-1}$ | [54] |
| $D_{NO_3^-}$ | $5.08 \times 10^{-12} T$ | m$^2$ s$^{-1}$ | [49] |
| $D_{CO_2}$ | $14.6836 \times 10^{-9}\left(\frac{T}{217.2056} - 1\right)^{1.997}$ | m$^2$ s$^{-1}$ | [54] |
| $D_{O_2}$ | $14.6836 \times 10^{-9}\left(\frac{T}{217.2056} - 1\right)^{1.997}$ | m$^2$ s$^{-1}$ | [54] |
| *Acid-base reactions* | | | |
| $S_1$ | -96.31 | J mol$^{-1}$ K$^{-1}$ | [57] |
| $S_2$ | -148.1 | J mol$^{-1}$ K$^{-1}$ | [57] |
| $S_W$ | -80.66 | J mol$^{-1}$ K$^{-1}$ | [57] |
| $H_1$ | 7.64 | kJ mol$^{-1}$ | [57] |
| $H_2$ | 14.85 | kJ mol$^{-1}$ | [57] |
| $H_W$ | 55.84 | kJ mol$^{-1}$ | [57] |
| $k_1$ | $\exp\left(1246.98 - \left(6 \times \frac{10^4}{T} - 183 \ln(T)\right)\right.$ | s$^{-1}$ | [60] |
| $k_2$ | 59.44 | s$^{-1}$ | [60] |
| $k_3$ | 2.23×10$^3$ | L mol$^{-1}$ s$^{-1}$ | [60] |
| $k_4$ | 6.0×10$^9$ | L mol$^{-1}$ s$^{-1}$ | [60] |
| $k_W$ | 2.4×10$^{-5}$ | L mol$^{-1}$ s$^{-1}$ | [61] |

| | | | |
|---|---|---|---|
| *Gas solubility* | | | |
| $A_1$ | -60.2409 | -- | 62 |
| $A_2$ | 93.4517 | -- | 62 |
| $A_3$ | 23.3585 | -- | 62 |
| $B_1$ | $2.3517\times10^{-2}$ | -- | 62 |
| $B_2$ | $-2.3656\times10^{-2}$ | -- | 62 |
| $B_3$ | $4.7036\times10^{-3}$ | -- | 62 |
| *Electrode reactions – anode* | | | |
| $i_{0,\mathrm{OER}}$ | $1\times10^{-8}$ | A $cm^{-2}$ | 63 |
| $E^0_{\mathrm{OER}}$ | 1.23 | V | 61 |
| $\alpha_{\mathrm{a,OER}}$ | 1.7 | -- | 63 |
| $\alpha_{\mathrm{c,OER}}$ | 0.1 | -- | 63 |
| *Electrode reactions – biocatalyst layer* | | | |
| $i_{0,\mathrm{X}}$ | $1.368\times10^{-8}$ | A $cm^{-2}$ | 50 |
| $E^0_{\mathrm{X}}$ | 0.314 ($QH_2$)<br>0.064 (NADH) | V | Eq. 7, 8 |
| $\alpha_{\mathrm{a,X}}$ | 0.5 | -- | 50 |
| $\alpha_{\mathrm{c,X}}$ | 0.5 | -- | 50 |
| *Electron transport* | | | |
| $\sigma_{\mathrm{GDE}}$ | 220 | S $m^{-1}$ | 42 |

*Supplementary Note 1: Intracellular $CO_2$-fixation enzyme concentration*

To estimate the intracellular concentration of formate dehydrogenase (which we assume to be the rate-limiting step in the reductive glycine pathway, following Bar Even *et al.*[58]), we use the estimate that the total intracellular protein count is ~$2.36 \times 10^6$,[45] and assume that formate dehydrogenase comprises ~3% of the total protein count. This number is based on the estimate that RuBisCo comprises ~3% of all proteins in autotrophic microbes that use the Calvin cycle to fix carbon.[44] We also assume a cell diameter of 1 µm, in accordance with the geometric assumptions regarding the bCL.

10 D. Gupta, M. C. Sutherland, K. Rengasamy, J. Mark Meacham, R. G. Kranz and A. Bose, *MBio*,

2019, **10**, 1659–1677.
11 V. Agostino and M. A. Rosenbaum, *Front. Energy Res.*, 2018, **6**, 1–10.
12 A. R. Rowe, S. Xu, E. Gardel, A. Bose, P. Girguis, J. P. Amend and M. Y. El-Naggar, *MBio*, 2019, **10**, 1659–1677.
13 S. T. Lohner, J. S. Deutzmann, B. E. Logan, J. Leigh and A. M. Spormann, *ISME J.*, 2014, **8**, 1673–1681.
14 P.-L. Tremblay, N. Faraghiparapari and T. Zhang, *Catalysts*, 2019, **9**, 166.
15 M. Firer-Sherwood, G. S. Pulcu and S. J. Elliott, *J. Biol. Inorg. Chem.*, 2008, **13**, 849–854.
16 K. Schuchmann and V. Müller, *Nat. Rev. Microbiol.*, 2014, **12**, 809–821.
17 P. L. Tremblay, T. Zhang, S. A. Dar, C. Leang and D. R. Lovley, *MBio*, , DOI:10.1128/mBio.00406-12.
18 B. Kraft, M. Strous and H. E. Tegetmeyer, *J. Biotechnol.*, 2011, **155**, 104–117.
19 C. Moreno-Vivián, P. Cabello, M. Martínez-Luque, R. Blasco and F. Castillo, *J. Bacteriol.*, 1999, **181**, 6573–6584.
20 A. Tiemeyer, H. Link and D. Weuster-Botz, *Appl. Microbiol. Biotechnol.*, 2007, **76**, 75–81.
21 M. D. Youngblut, O. Wang, T. P. Barnum and J. D. Coates, *Annu. Rev. Microbiol.*, 2016, 70, 435–457.
22 J. D. Coates and L. A. Achenbach, *Nat. Rev. Microbiol.*, 2004, **2**, 569–580.
23 L. L. Barton and G. D. Fauque, in *Advances in Applied Microbiology*, Elsevier Inc., 1st edn., 2009, vol. 68, pp. 41–98.
24 A. A. Santos, S. S. Venceslau, F. Grein, W. D. Leavitt, C. Dahl, D. T. Johnston and I. A. C. Pereira, *Science (80-. ).*, 2015, **350**, 1541–1545.
25 C. Rückert, *Curr. Opin. Microbiol.*, 2016, **33**, 140–146.
26 N. J. Claassens, C. A. R. Cotton, D. Kopljar and A. Bar-Even, *Nat. Catal.*, 2019, **2**, 437–447.
27 A. J. Abel, J. D. Adams and D. S. Clark, *bioRxiv*, 2021, 2021.07.01.450744.
28 International organization for standarization, *ISO 14040*.
29 International Organization for Standardization (ISO), *ISO 14044*.
30 D. M. de S. Simone Manfredi, Karen Allacker, Kirana Chomkhamsri, Nathan Pelletier, *Product Environmental Footprint (PEF) Guide*, 2012.
31 J. Huang, B. Mendoza, J. S. Daniel, C. J. Nielsen, L. Rotstayn and O. Wild, *Clim. Chang. 2013 Phys. Sci. Basis Work. Gr. I Contrib. to Fifth Assess. Rep. Intergov. Panel Clim. Chang.*, 2013, **9781107057**, 659–740.
32 J. G. Chen, R. M. Crooks, L. C. Seefeldt, K. L. Bren, R. M. Bullock, M. Y. Darensbourg, P. L. Holland, B. Hoffman, M. J. Janik, A. K. Jones, M. G. Kanatzidis, P. King, K. M. Lancaster, S. V Lymar, P. Pfromm, W. F. Schneider and R. R. Schrock, *Science (80-. ).*, 2018, **360**, eaar6611.
33 V. Singh, I. Dincer and M. A. Rosen, eds. I. Dincer, C. O. Colpan and O. B. T.-E. Kizilkan Energetic and Environmental Dimensions, Academic Press, 2018, pp. 935–959.
34 United State Environmental Protection Agency (EPA), *Off. Air Radiat.*, 2010, 1–43.
35 C. T. Corporation, *Final Report on SURVEY OF ELECTROCHEMICAL PRODUCTION OF INORGANIC COMPOUNDS*, Argonne, IL, 1980.
36 A. Al-Dallal, *Al-Khwarizmi Eng. J.*
37 I. Garcia-Herrero, M. Margallo, R. Onandía, R. Aldaco and A. Irabien, *Sci. Total Environ.*, 2017, **580**, 147–157.
38 S. Deutz and A. Bardow, *Nat. Energy*, 2021, **6**, 203–213.
39 R. Stropnik, A. Lotrič, A. Bernad Montenegro, M. Sekavčnik and M. Mori, *Energy Sci. Eng.*, 2019, **7**, 2519–2539.
40 J. Xu, G. Liu, J. Li and X. Wang, *Electrochim. Acta*, 2012, **59**, 105–112.
41 J. Izursa, E. A. Hanlon, N. Y. Amponsah and J. C. Capece, *Carbon Footprint of Biofuel Sugarcane Produced in Mineral and Organic Soils in Florida - Manuscript submitted for publication*, LaBelle,

FL, 2013.
42 L. C. Weng, A. T. Bell and A. Z. Weber, *Phys. Chem. Chem. Phys.*, 2018, **20**, 16973–16984.
43 N. J. Claassens, G. Bordanaba-Florit, C. A. R. Cotton, A. De Maria, M. Finger-Bou, L. Friedeheim, N. Giner-Laguarda, M. Munar-Palmer, W. Newell, G. Scarinci, J. Verbunt, S. T. de Vries, S. Yilmaz and A. Bar-Even, *Metab. Eng.*, 2020, **62**, 30–41.
44 Y. M. Bar-On and R. Milo, *Proc. Natl. Acad. Sci. U. S. A.*, 2019, **116**, 4738–4743.
45 R. Milo, *BioEssays*, 2013, **35**, 1050–1055.
46 K. B. Andersen and K. Von Meyenburg, *J. Bacteriol.*, 1980, **144**, 114–123.
47 A. Sydow, T. Krieg, R. Ulber and D. Holtmann, *Eng. Life Sci.*, 2017, **17**, 781–791.
48 F. Salimijazi, J. Kim, A. M. Schmitz, R. Grenville, A. Bocarsly and B. Barstow, *Joule*, 2020, **4**, 2101–2130.
49 A. J. Abel and D. S. Clark, *ChemSusChem*, 2021, **14**, 344–355.
50 A. J. Abel, J. M. Hilzinger, A. P. Arkin and D. S. Clark, *Bioelectrochemistry*, 2022, 108054.
51 M. Lin, L. Han, M. R. Singh and C. Xiang, *ACS Appl. Energy Mater.*, 2019, **2**, 5843–5850.
52 M. R. Singh, E. L. Clark and A. T. Bell, *Phys. Chem. Chem. Phys.*, 2015, **17**, 18924–18936.
53 Fuel Cell Store, *Sigracet GDL 34 & 35 Series Gas Diffusion Layer*, 2020.
54 R. E. Zeebe, *Geochim. Cosmochim. Acta*, 2011, **75**, 2483–2498.
55 P. Han and D. M. Bartels, *J. Phys. Chem.*, 1996, **100**, 5597–5602.
56 E. L. Cussler, *Diffusion: Mass Transfer in Fluid Systens*, Cambridge University Press, Cambridgeshire, New York, 3rd edn., 2009.
57 D. R. Lide, Ed., *CRC Handbook of Chemistry and Physics, 84th edition*, CRC Press, 84th edn., 2004.
58 A. Bar-Even, E. Noor, A. Flamholz and R. Milo, *Biochim. Biophys. Acta - Bioenerg.*, 2013, **1827**, 1039–1047.
59 D. A. Stolpera, N. P. Revsbech and D. E. Canfield, *Proc. Natl. Acad. Sci. U. S. A.*, 2010, **107**, 18755–18760.
60 K. G. Schulz, U. Riebesell, B. Rost, S. Thoms and R. E. Zeebe, *Mar. Chem.*, 2006, **100**, 53–65.
61 P. W. Atkins, *Physical Chemistry*, Oxford University Press, 4th edn., 1990.
62 Ulf Riebesell, Victoria J. Fabry, Lina Hansson and Jean-Pierre Gattuso, *Guide to best practices for ocean acidification research and data reporting*, 2011.
63 S. Haussener, C. Xiang, J. M. Spurgeon, S. Ardo, N. S. Lewis and A. Z. Weber, *Energy Environ. Sci.*, 2012, **5**, 9922–9935.
